# FourC: identifying significant and differential contacts in 1D chromatin conformation data

**DOI:** 10.64898/2026.03.05.709811

**Authors:** Wilfred Wong, Samuel J Kaplan, Renhe Luo, Julian Pulecio, Jielin Yan, Danwei Huangfu, Christina S. Leslie

## Abstract

4C-seq is a cost-effective 3C-based assay that measures the interactions between a single genomic element and all other genomic elements. However, 4C-seq data remains semi-quantitative because it cannot be deduplicated without UMIs. To address this, we developed an open source method, FourC, based on a Bayesian Bernoulli regression model, that overcomes the duplication problem and models spatial patterns with Gaussian processes to identify significantly enriched and differential contacts. We demonstrate the utility of FourC on 4C-seq data that profiles the local chromatin structure at key genes necessary for pancreatic differentiation and under CRISPR perturbation of enhancers.

## Background

4C-seq measures the spatial proximity between one genomic element, called the bait, and all other genomic elements, called the captures. These data give insight into regulatory enhancer-promoter interactions, as well as structural interactions that shape the genome. During this procedure, spatially proximal bait-capture pairs are first converted into circularized fragments under restriction digest conditions before being subjected to PCR and sequencing. Two measurement issues render the resulting data semi-quantitative: i) PCR duplication effects; and ii) 3C sampling biases. 4C-seq counts cannot be positionally deduplicated due to the deterministic restriction digest step (although this limitation can be overcome with an additional sonication step); thus distinct interaction counts cannot be distinguished from PCR products. Moreover, 4C-seq is affected by 3C sampling biases, such as those attributable to fragment-level GC content, length, and distance from the bait. Previous analysis strategies have indirectly addressed these issues, but tend to involve averaging or smoothing over genomic windows, which simultaneously reduces the resolution of the assay and introduces an additional analysis parameter [1–10].

These measurement issues can make it challenging to identify significant interactions within experiments and differential interactions between experiments, since the read counts are partially a function of the PCR step. To overcome these issues, we establish a relationship between the 4C-seq reaction products and the observed read counts to motivate the transformation of fragment counts into binary presence/absence indicators and overcome the duplication problem [1, 11]. These developments lead to a Bernoulli spatial generalized linear mixed model (SGLMM), which allows us to condition on the remaining 3C measurement effects and estimate fragment-level contact frequencies. Our model can then be used for: i) the identification of significantly enriched contacts within a 4C-seq experiment; and ii) the detection of differential contacts between 4C-seq experiments. To make the model accessible, we utilize the Integrated Nested Laplace Approximation (INLA; 12) and provide the model, along with helpful utilities, in the FourC package (Figure 1).

**Figure 1:**
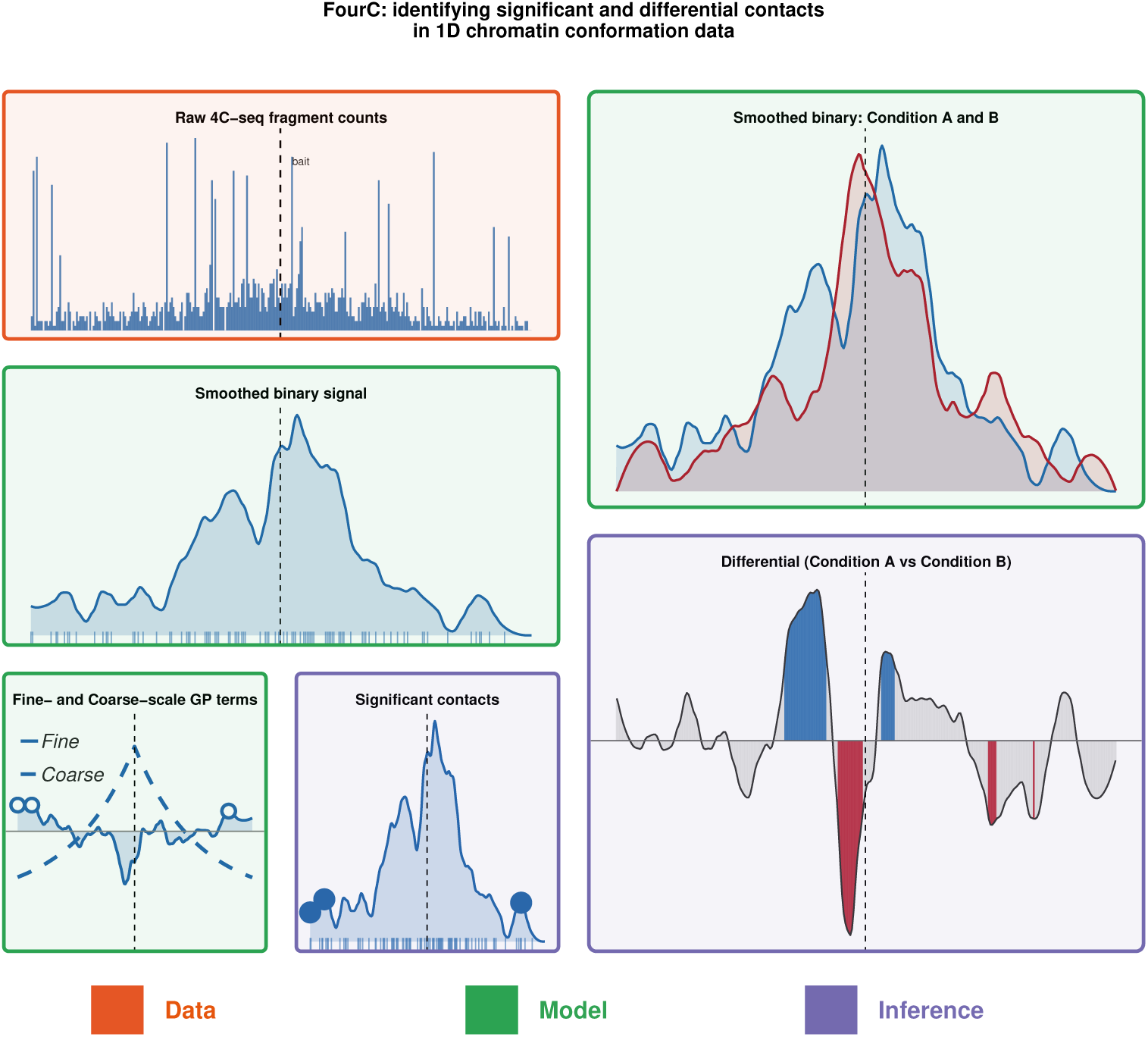
FourC model. 4C-seq counts are binarized to overcome PCR duplication, then decomposed into fine- and coarse-scale spatial signals used to nominate loops and differential interactions. Opaque color boxes mark the relevance of the data, the model, and the inferences.

To demonstrate the utility of our method we applied our FourC methodology to understand the evolution of enhancer-promoter contacts during differentiation and under enhancer perturbations in a human pancreatic stem cell system where embryonic stem cells (ES) differentiate stepwise through definitive endoderm (DE), primitive gut tube (FG), and pancreatic progenitor (PP) states. 4C-seq data collected under wild-type (WT) conditions for *GATA6, PDX1, ONECUT1*, and *SOX17* were used to identify distal regulatory elements that establish significant long-range contacts with the promoter of these genes during differentiation [13–15]. Next, for *GATA6* and *ONECUT1*, we measured chromatin topology prior to and under CRISPR-based silencing or deletion of their enhancers. These analyses showed that for some key developmental genes, enhancer-promoter interactions are established at early stages of differentiation, before activation of the target gene. We also confirmed that CRISPRi-based enhancer perturbations have a weaker effect on promoter interactions compared with CRISPR-Cas9 enhancer deletion.

## Results

### Motivation for a binarized fragment model

To illustrate the problem with duplicated fragment counts and motivate binarization, we will use 4C-seq data from the *ONECUT1* promoter at the PP stage. The replicate-level data, centered on a 2.4 Mb window around the bait and conditioned on the blind vs. non-blind fragmentation class (see [3] for a definition), exhibit a partial decay in fragment counts, but large counts ∼1 Mb away are likely attributable to PCR artifacts. These issues cannot be remedied by window-based smoothing (Figure 2a) and this increases replicate-to-replicate variation. Matched Hi-C data suggests that the observed counts at those distances are likely more attributable to PCR effects than contacts, since those fairly large signals are unobservable in Hi-C (Figure 2c).

**Figure 2:**
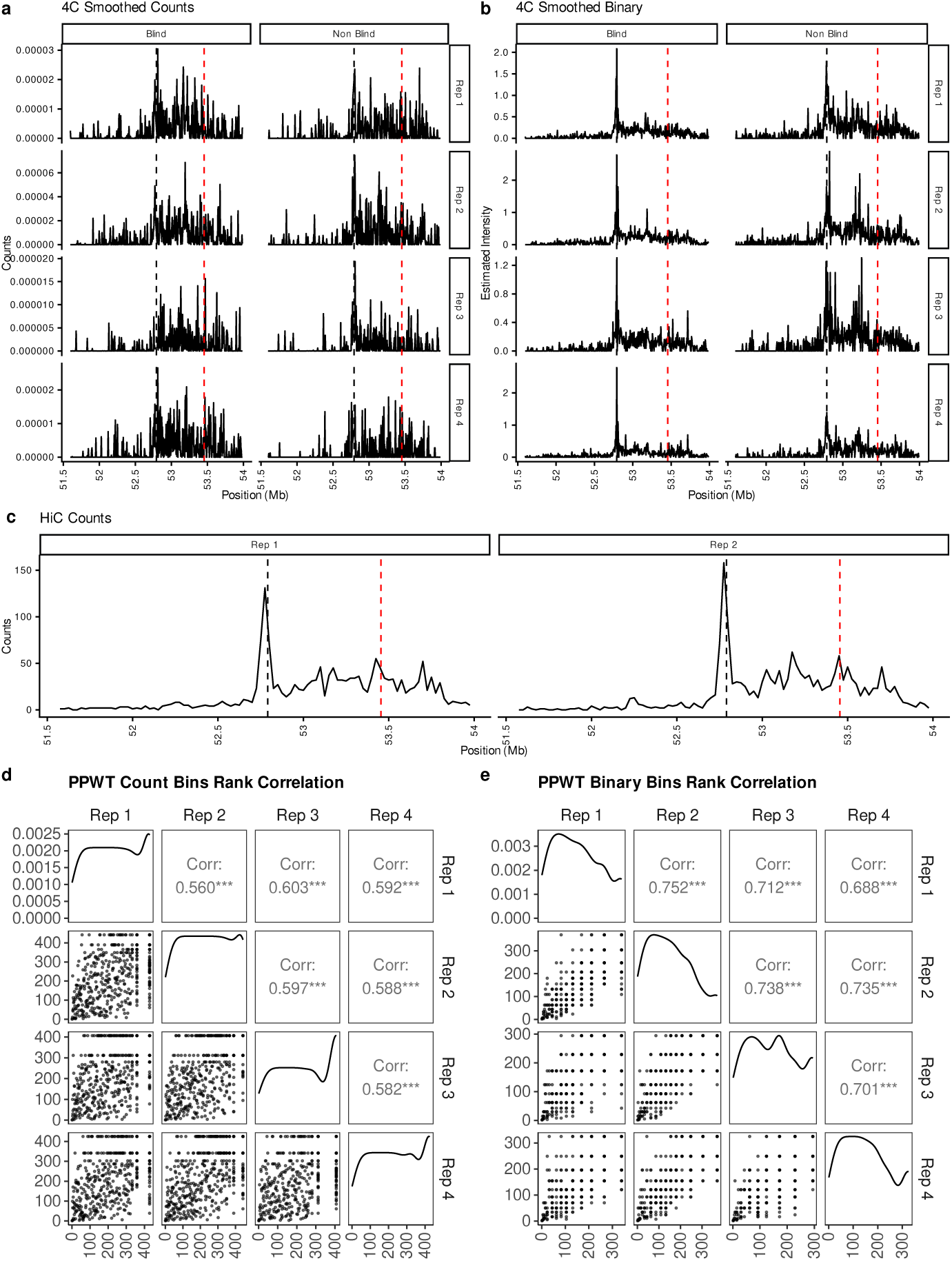
Binarization improves reproducibility between replicates. **(a)** A 2.4 Mb window around the *ONECUT1* viewpoint, binned into 5 kb bins with summed counts, split by blind and non-blind fragments. Dashed line: enhancer +e664 (Kaplan et al.). Black line: viewpoint of an experiment. **(b)** As in (a), but counts are first binarized and converted to a capture intensity via cloglog transform. **(c)** Hi-C counts for the same region around the *ONECUT1* promoter. **(d)** Binned count correlations across four replicates, compared to binned capture-probability correlations in **(e)**.

In contrast, a smoothed binary estimator, which is a function of the capture rate (see Methods), removes the artifactual contacts at ∼1 Mb away (Figure 2b). Furthermore, the contact profile suggests a more typical monotonic decay in contacts and higher concordance between replicates. The pairwise rank concordance for the count data (Figure 2d) is improved by binarization (Figure 2e) as a consequence of removing the variation likely attributable to PCR duplication (see Fig. S6 for additional examples).

### A statistical model for binarized fragment data

In the 4C-seq procedure, we model the initial count of the *i*th fragment prior to PCR as *y_i_* ∼ Pois(*λ_i_*), where *λ_i_* (capture rate) depends on spatial proximity (contact frequency) and 3C parameters [16]. In a Galton-Watson branching process (see Supplemental Note), the duplicated fragment count *ρ_i_*(*y_i_, p_d,i_, c*) depends on the starting count, *y_i_*, amplification efficiency, *p_d,i_*, and PCR cycles, *c*. Read counts are then *r_i_*|*ρ_i_* ∼ Pois(*Rρ_i_*), where *R* = *N_R_/* Σ*_j_ ρ_j_* is the fraction of molecules sampled at depth *N_R_*.

We therefore propose a binarized formulation of the problem. Initially, this seems *ad hoc* because it is unclear how a *capture probability*, the probability of observing fragment *i* at least once after sequencing, is related to its intrinsic *capture rate λ_i_*. Initial attempts at binarizing the observations were met with resistance, owing to the intuition that information is lost; while some information is indeed discarded, we sketch here why this is advantageous [1].

Assuming that PCR is a largely lossless process and that libraries are sufficiently sequenced, so that ℙ(*r_i_* = 0|*y_i_ >* 0) ≈ 0, then a fragment can only be observed if *y_i_ >* 0 (Fig. S7 demonstrates that most libraries are saturated in practice [17]). Hence, even if *r_i_* depends on PCR duplication, the presence/absence indicators reveal information about the original data before PCR. Under these assumptions, a *capture probability* is fundamentally tied to the intrinsic *capture rate λ*, since ℙ(*r_i_* = 0) ≈ ℙ(*y_i_* = 0) = *e*^−^*^λi^* . A more careful argument is given in the Supplemental Note.

Defining *o_i_*= 1[*r_i_ >* 0], then *o_i_*∼ *Ber*(*p_i_*)*, p_i_* = 1 − *e*^−*λi*^ , and through the complementary log-log transform, we may perform estimation and inference for *λ_i_*without considering the effects of PCR [18, 19].

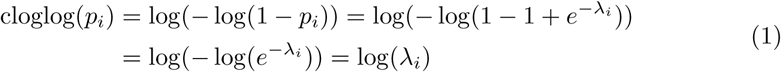

As the data arises at the fragment level, and the libraries are readily saturated, we may model at the fragment level rather than binning. For a collection of experiments collected from the same bait, let *o_ij_* be the binary capture indicator, where *i* is the *i*th fragment in the region under consideration from the *j*th replicate. Each measurement is associated with technical covariates, *x_ij_*, and the position of the fragment *s_ij_*. Our data thus consists of (*o_ij_, x_ij_, s_ij_*), for *i* ∈ [1*, …, F*]*, j* ∈ [1*, …, J*], where we have a total of *F* fragments, up to say ±1.2 Mb, around the bait and *J* replicates in the experiment.

We assume a latent Gaussian model (LGM), where the observation likelihood is linked to structural predictors that are assumed to have a Gaussian form [12]. The fixed effects consist of fragment-level characteristics, such as GC, length, type (blind/non-blind), and sample indicators, which adjust for differences in starting cell material and overall reaction efficiency for an experiment. The effect of fragment length and GC content is modeled with natural splines, denoted as f*_Length_* and f*_GC_* and can be used to flexibly model covariate effects (Figures 3a, 3b, 3c, 3d).

**Figure 3:**
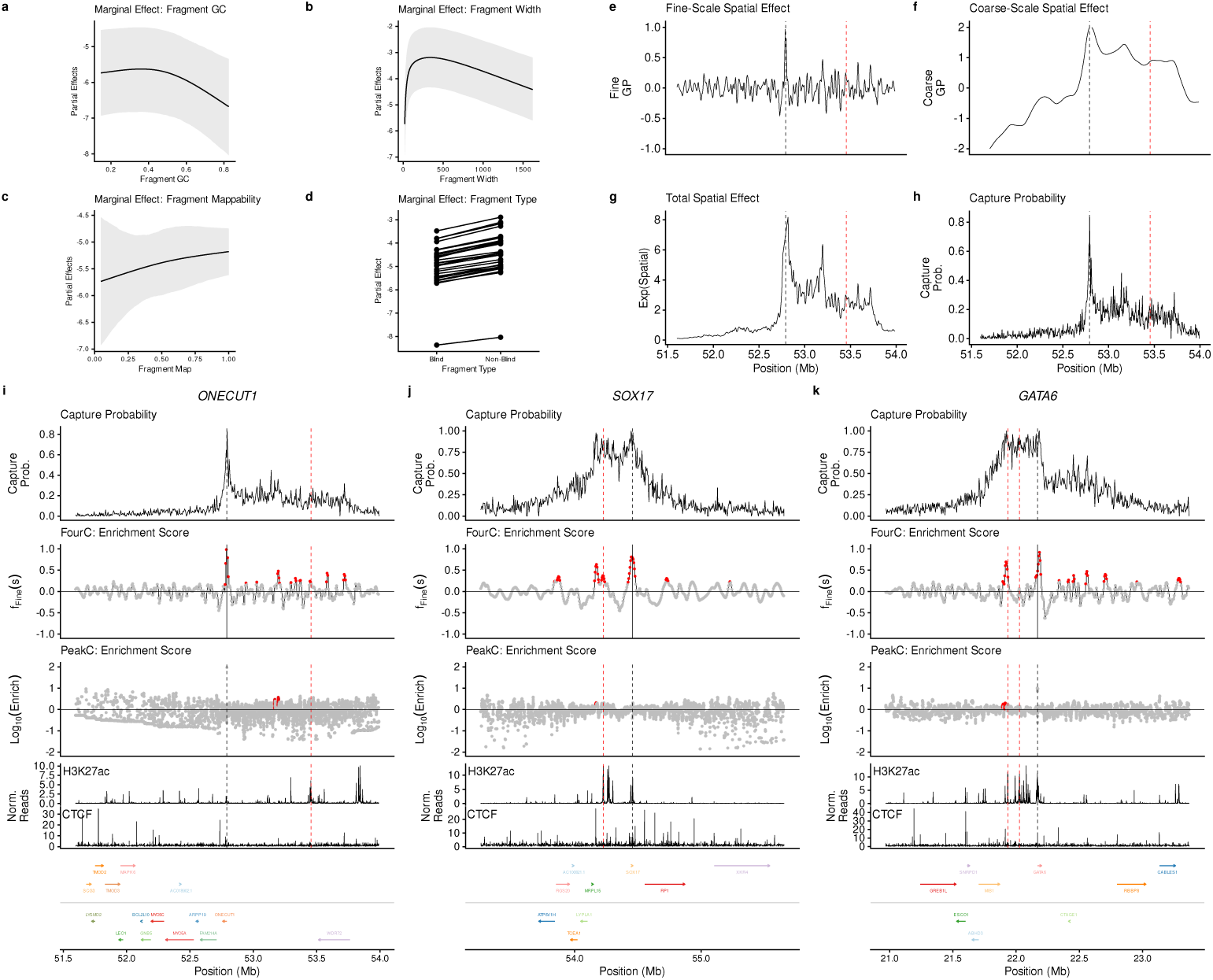
FourC modeling captures technical effects and spatially structured signals. **(a–d)** Average effects of fragment-level GC content, fragment length, mappability, and fragmentation class. **(e, f)** Fine- and coarse-scale Gaussian Process terms for *ONECUT1*. **(g)** Spatial terms summed and exponentiated to estimate capture intensity. **(h)** Capture probability, averaged across four *ONECUT1* replicates, compared to the estimate in **(g)**. **(i–k)** Example experiments for *ONECUT1* (PP), *SOX17* (DE), and *GATA6* (DE), with capture probabilities as above. Fine-scale term (FourC) and enrichment scores (peakC) are plotted, with red dots marking significant regions from each method; H3K27ac and CTCF are shown for context.

Next, our model includes two spatial (Matern Gaussian process) terms, *f ^Coarse^* to model large-scale (∼ 100 kb) smooth patterns, such as the overall distance decay, and *f ^F ine^* to capture residual spatial variation at a smaller scale (∼ 10 kb) [20, 21]. A computationally efficient approximation is used by relating a realization of a Matern process to the stationary solutions of a Stochastic Partial Differential Equation (SPDE) and this involves a discrete grid approximation (see Methods for further details). An example of the spatially smooth terms is shown for *ONECUT1* (Figures 3e, 3f) and the combined spatial terms (Figure 3g) can be compared to the capture probabilities (Figure 3h). The LGM is specified as follows:

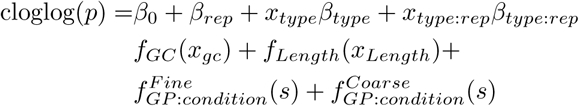

The spatial terms are allowed to vary between different logical groupings, such as experimental conditions. Furthermore, since the blind fragments are dependent on the rate at which non-blind fragments are formed, we will let these quantities vary between experimental samples. For all fixed effects of the model, we apply flat priors, and for the spatial random effects, we utilize penalized complexity priors (see Methods for further details).

### Identifying significant interactions and differential interactions

#### Identifying enriched contacts with fine-scale spatial terms

We apply FourC to identify significant interactions from 4C-seq experiments performed in a guided pancreatic differentiation system at the loci of key lineage-determining transcription factors: *ONECUT1*, *SOX17*, and *GATA6*. To examine the degree to which FourC improves over existing methods for prioritizing functional genomic regions, we evaluated peakC, r3Cseq, and FourSig.

In brief, peakC estimates the significance of 4C contacts by first using monotonic regression to estimate background contact frequencies across replicates, and then utilizing rank-product statistics to assign significance [5]. FourSig constructs a permutation null by shuffling the fragment positions and examining the background distribution of read counts within a fixed window [1]. Note that this approach does not sufficiently account for genomic distance biases. r3Cseq uses a smoothing spline to estimate the background contact rate prior to z-scoring and testing against a standard normal null [22].

We find that FourSig and r3Cseq struggle with the specific biases of these datasets. While FourSig identifies many true functional genomic elements, it also tends to identify numerous regions that are not aligned with orthogonal epigenomic data. At an FDR of 0.1, FourSig essentially identifies any region that passes a raw count threshold (Fig. S5), indicating low specificity. Conversely, r3Cseq does not report any significant events across all experiments considered in this setting (beyond the bait-proximal fragment). Because its smoothing spline is unconstrained, it can yield negative expected values, rendering the subsequent z-scoring procedure overly conservative even though it attempts to account for genomic distance bias. Consequently, we compare primarily against peakC, which is the most well-maintained and robust method for analyzing 4C-seq data [5].

Our SGLMM contains a coarse-grained and fine-grained spatial intercept term, and this allows us to condition on large-scale spatial patterns (coarse) and identify local enrichments (fine) in a manner analogous to existing 3C methods for identifying significant contacts [20, 21]. Operationally, this corresponds to using the posterior distribution, p(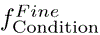 (s*_i_*)|y), and finding regions where p(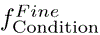 (s*_i_*) > 0|y) > 1 − *α*, for some decision threshold *α*. As our inferences depend on the random effects of our model, we assessed the sensitivity of our inferences to the mesh built on top of the spatial domain and the choice of hyperpriors and found that our inferences were largely insensitive to these choices (see Fig. S8). Marginal statements are typically of interest, but joint probability statements can be obtained through excursion sets [23].

Our proposed methodology—namely, finding regions where the probability of the posterior excluding 0 is at least 90% (see Methods, Supplementary Note)—can identify recently discovered functional enhancers, which are marked by red vertical lines. *ONECUT1* ‘s functional enhancer is flagged by our method, and the other regions that are flagged are enriched for CTCF binding (Figure 3i). In contrast, peakC does not identify this enhancer at an FDR of 0.1. In the case of *SOX17*, the functional enhancer, SOX17+e10, is flagged by our method, but not by peakC, and additionally, we find many more CTCF and H3K27ac enrichments (Figure 3j). Finally, in the case of *GATA6*, only one of two enhancers, to be discussed more later and denoted as +e12 and +e9 (the leftmost and rightmost red dashed lines respectively), is found by our proposed method, but neither of them is found by peakC (Figure 3k). As shown by these examples, FourC identifies significant contacts that are functionally relevant while eliminating duplication effects and conditioning on technical covariates. Fig. S9 and Fig. S10 provide quantitative summaries and permutation controls. Additional called peaks are provided in Fig. S14, Fig. S16, Fig. S15, and Fig. S17.

#### Identifying differential contacts with total spatial terms

Next, we applied FourC to obtain log fold changes for *GATA6*, *ONECUT1*, and *SOX17* between the DE and ES stages, and for *PDX1* between the PP1 and ES stages; extant Hi-C data are available to validate log fold changes for all four comparisons. While 4C-seq and Hi-C utilize different digestion patterns, these technical effects are constant within each experiment; consequently, these technical effects should cancel out during the calculation of fold changes. Therefore, we anticipated that our fold changes from FourC should match those of deduplicated Hi-C data.

Our SGLMM contains the elements to compute log fold changes that do not depend on binning or peak calling. For conditions A and B, we define a continuous fold change, which can be binned for comparison purposes by aggregating the posteriors over the fragments into windows:

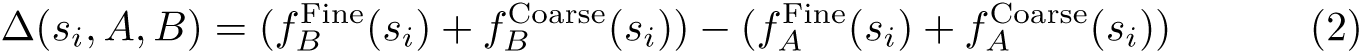

The log fold changes obtained through FourC are compared to those obtained by DESeq2, and in general, the raw log fold changes tend to be shrunk to zero relative to the DESeq2 estimates as a consequence of the spatial prior (Figure 4a). Across multiple experimental comparisons with matching Hi-C stages for validating the comparisons, the RMSE of the log fold changes obtained by FourC is smaller than those obtained by DESeq2 when compared against the Hi-C log fold change estimates (Figure 4b). These metrics were stratified because distal captures occur less frequently and are more difficult to model.

**Figure 4:**
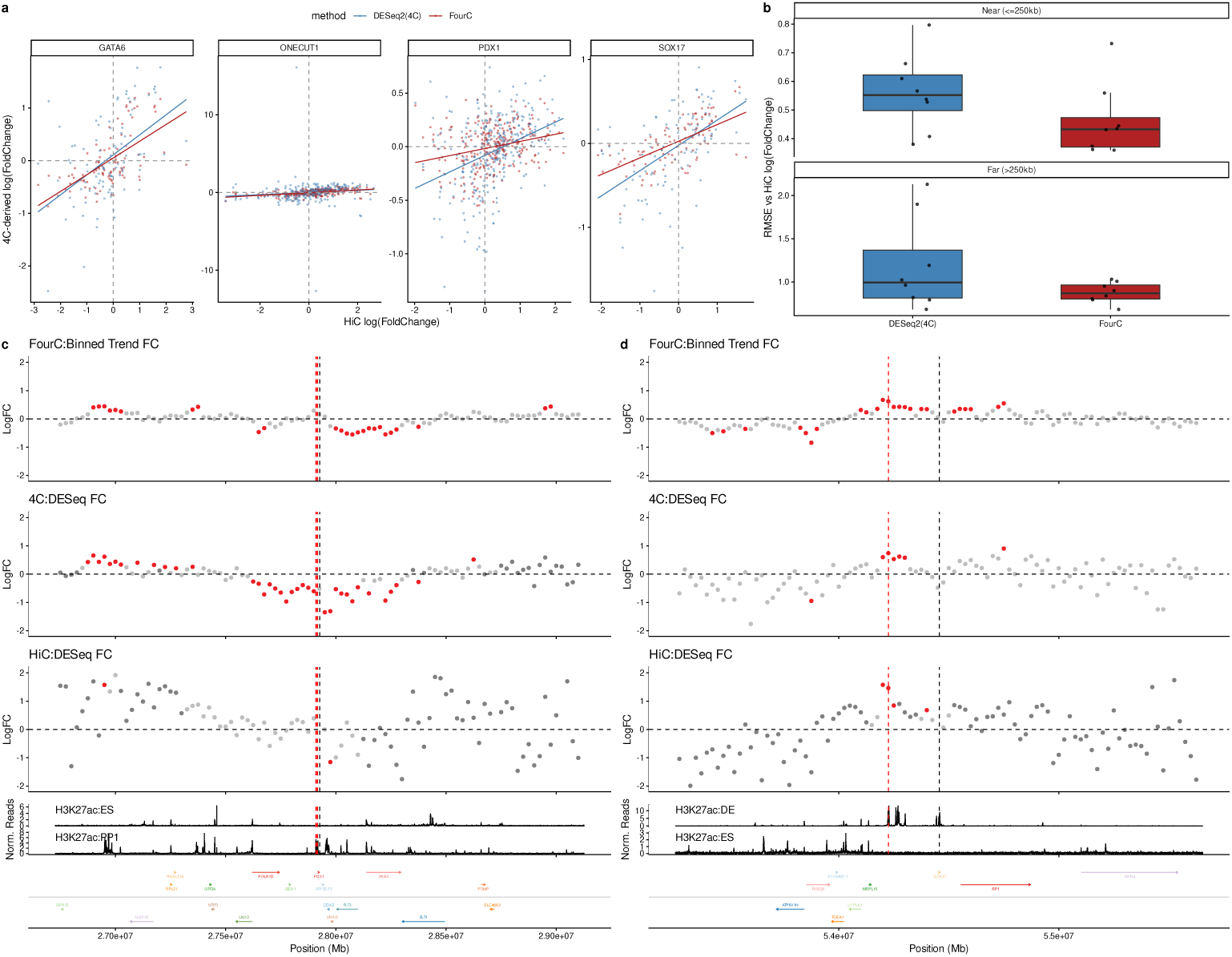
FourC estimates log fold changes more accurately. **(a)** 4C log fold changes vs. Hi-C at the DE stage; DESeq2 (blue) vs. FourC (red). **(b)** Log fold changes from *GATA6*, *SOX17*, *ONECUT1*, and *PDX1* (differentials between consecutive stages of ES, DE, PP) compared to matched Hi-C, with RMSE stratified by distance (distal estimates are noisier). **(c, d)** FourC and DESeq2 log fold changes for *PDX1* (PP vs. ES) and *SOX17* (DE vs. ES) across differentiation, compared to Hi-C-derived DESeq2 estimates.

Significant differential interactions occur when the central credible interval excludes zero with high probability: 1[0 ∈ C*_α_*(Δ(*s_i_, A, B*))]. Crucially, this approach leverages information between spatially adjacent regions of the genome to inform our estimates and inferences, providing a more robust estimate of chromatin changes than fragment-wise comparisons (see Methods that relate Δ to fold changes).

Notably, fold change estimates derived from 4C read counts (labeled as ‘4C:DEseq FC’) disagree at the enhancer region. DESeq applied to the 4C counts suggests that *PDX1* induction is associated with a loss of contact against a promoter-proximal enhancer, while FourC is in agreement with the Hi-C data (Figure 4c). Such disagreement may be due to PCR duplication effects near the viewpoint. Outside of the viewpoint region, the experiments all appear to flag an upstream region that is marked by histone acetylation.

Next, we examined *SOX17* during the differentiation towards the DE state (Figure 4d). A functional enhancer element, *SOX17* +e10, has been identified upstream of the gene promoter approximately 200 kb away [13]. All methods indicate an increase in enhancer-promoter contacts during lineage specification. Note, this enhancer would have been missed if we pre-specified a loop atlas with peakC, so standard methods based on calling loops prior to testing for differences can fail to detect important functional elements. Therefore, analyzing the entire genomic window around the bait for changes is a more robust approach. Further, we see that the aggregation of the fragment-level posteriors results in more resolution consistent inferences (see Fig. S12, Fig. S11).

### Examining chromatin topology under enhancer perturbations

While our previous differentiation experiments demonstrate that enhancer-promoter contacts accumulate and coincide with gene activation, the relationship between looping and enhancer function remains unclear. It is possible that the accumulation of contacts is driven by the activity of the enhancer; alternatively, the accumulation of contacts can occur in a manner independent of the enhancer’s functional status.

To interrogate the dependency between loops and enhancer function, we applied our FourC method to study chromatin topology at *GATA6* and *ONECUT1*, two critical lineage-determining transcription factors for the definitive endoderm and pancreatic progenitor states, respectively, under CRISPR perturbations of their associated enhancer(s).

#### CRISPRi and CRISPR of *GATA6* enhancers during definitive endoderm formation

*GATA6* is a core definitive endoderm transcription factor that is required for proper definitive endoderm fate induction. Recent reports have identified two strong enhancers denoted *GATA6* +e9 and *GATA6* +e12 located in the TAD containing the gene which increase in contact with the promoter during differentiation, though the contacts appear to generically increase within the TAD [13]. Whether this accumulation of these contacts depends on a functional enhancer remains unknown.

To further parse the relationship between enhancer function and looping, we utilized both CRISPRi and CRISPR-Cas9 to target e9 and e12. While CRISPR-Cas9 physically deletes the underlying sequence, CRISPRi silences enhancer activity without affecting the underlying sequence. These two complementary approaches allowed us to examine how varying levels of enhancer perturbations can alter chromatin topology. Note that in both cases, the CRISPR enzyme and guide are introduced at the ES stage, so that the enhancer is perturbed throughout differentiation.

CRISPRi perturbations of +e9 and +e12 (the right and left red lines respectively) (Figure 5a, Figure 5b) show that the local chromatin interactions around +e9 and +e12 decrease, while the surrounding chromatin contacts are largely unchanged. These CRISPRi effects appear highly localized and subtle, such that an application of DESeq2 to the binned (25 kb) count data fails to identify those localized changes. This indicates that FourC is well suited for parsing subtle changes in genomic organization. Fig. S11 and Fig. S12 provide different bin sizes and a demonstration of resolution consistency.

**Figure 5:**
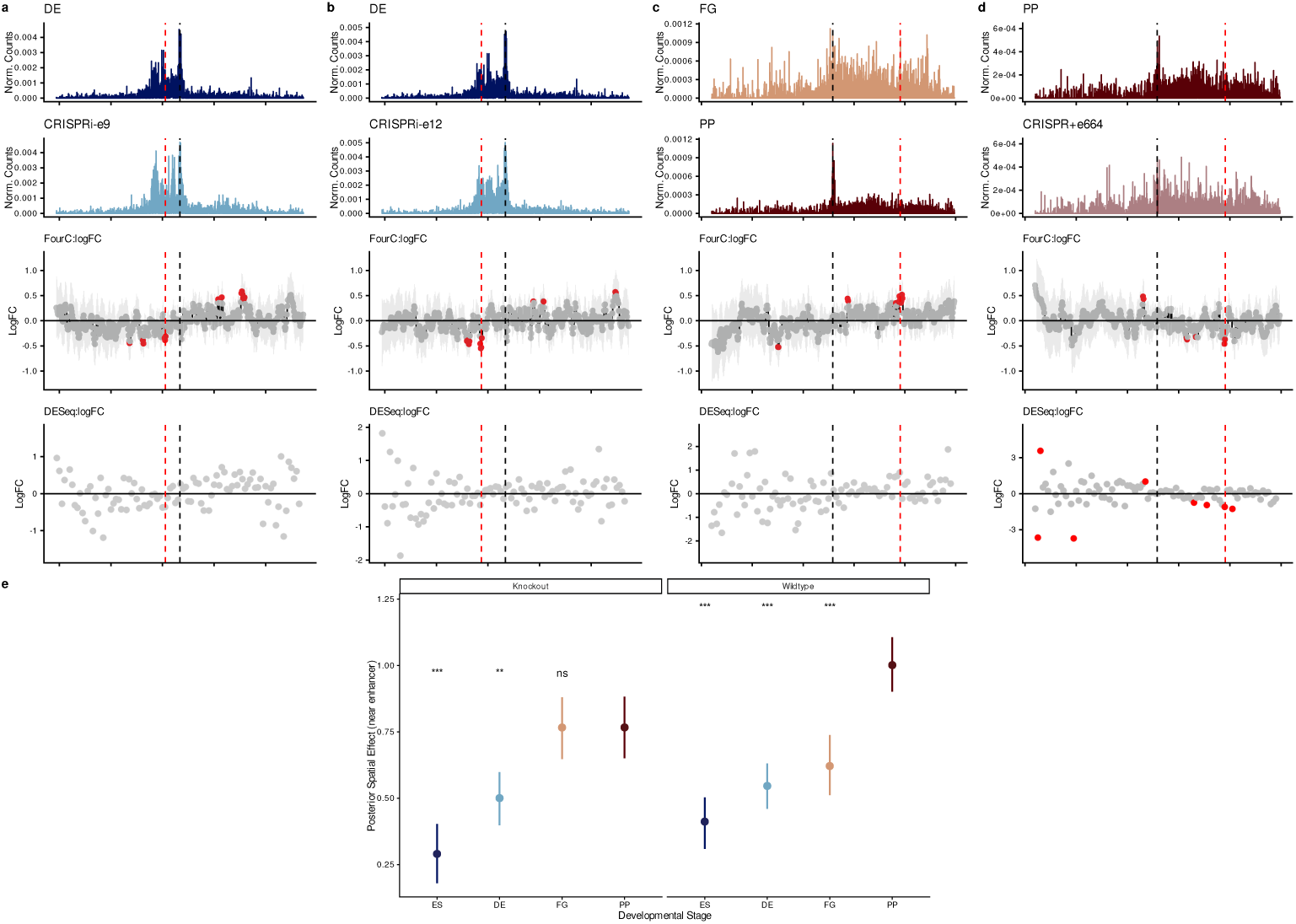
FourC dissects subtle effects of development and perturbation. **(a–d)** Effects of *GATA6* +CRISPRi-e9, *GATA6* +CRISPRi-e12, *ONECUT1* FG-to-PP differentiation, and *ONECUT1* +CRISPR+e664 at PP; FourC reliably detects perturbation effects across all four. **(e)** *ONECUT1* enhancer-promoter contacts evolve normally during differentiation but stall upon PP lineage specification.

Next, we performed CRISPR knockouts against +e9, +e12, and +e9/+e12 simultaneously (see Fig. S13). Subsequently, enhancer-promoter contacts drop against the targeted regions. This also leads to general decreases in interactions in the TAD (to the left of the promoter) containing *GATA6*, and increases in contact in the adjacent TAD (to the right of the viewpoint).

#### CRISPR perturbation of e664 at *ONECUT1* across differentiation

Our findings indicate that CRISPRi-based perturbations produce more subtle chromatin looping phenotypes than CRISPR-Cas9. In either case, loss of enhancer function directly reduces chromatin looping. However, since this experiment set assesses chromatin structure in the stage immediately after the ES state, we cannot determine if there was any accumulation of loops prior to DE. Thus, we turn our attention to *ONECUT1*, which is activated at the PP stage.

The enhancer of *ONECUT1*, called *ONECUT1+e664kb* and denoted as +e664 has recently been identified as a long-range enhancer 664 kb upstream of the promoter of *ONECUT1*. To examine how the contacts change during differentiation and under perturbation, we generated 4C-seq libraries at the ES, DE, FG, and PP stages under both wild-type and CRISPR-Cas9 settings. Upon induction of *ONECUT1*, the promoter of the gene increases in contact with +e664 between the DE and PP stages, and this coincides with the deposition of H3K27ac at +e664 [14]. Since we have collected FG data, we see that the EP interaction further strengthens from FG to PP, suggesting stepwise rewiring (Figure 5c). Upon knockout of +e664, the gene promoter loses contact with +e664 (Figure 5d), suggesting that the functional enhancer element mediates the looping of the promoter to +e664.

Finally, to assess the evolution of contacts around the enhancer in all stages, we examined the total spatial effects term of the model around +e664 in all four stages under WT and CRISPR-Cas9 settings. In both settings, the promoter of *ONECUT1* increases in contacts with +e664 between the ES and FG stage, suggesting that even in the absence of a functional enhancer, early establishment of these contacts is independent of the enhancer’s functional status (Figure 5e). However, in the PP stage, where *ONECUT1* is activated, contacts stall, suggesting that the increase in contacts in the wild-type setting is driven by a functional +e664.

Our analysis suggests a priming mechanism for the genome, whereby contacts are established in an enhancer-function independent manner during earlier cell states in order to facilitate the final commitment towards future cell lineages. This concept aligns well with the recently reported chromatin-competent regions, which are regions that lack epigenetic marks in the ES state, but are readily activatable via CRISPRa [15].

## Discussion

4C-seq is a methodology uniquely positioned for the high-resolution interrogation of chromatin topology, particularly for studying enhancer-promoter interactions under targeted genetic perturbations. While cell input and sequencing depth requirements are markedly lower than those of other 3C-based assays, the experimental design of 4C-seq inherently prevents positional deduplication. Although this limitation is frequently acknowledged in the literature, no analytical framework has yet been established to explicitly account for the resulting PCR effects at the fragment level.

Our model for the 4C-seq library assumes that the molecular products *y_i_* can be well modeled by a Poisson regression, but the observed read counts, *r_i_*, cannot due to PCR effects. Subsequently, we show that provided the zeros are by and large preserved during the sampling process, binarization arises as a natural solution for inferring the spatial proximity of DNA. In this framework, the presence or absence of a fragment—rather than its duplicated count—becomes the fundamental unit of information.

This conceptual advance leads to data transformations for visualization and a complementary log-log Bernoulli spatial generalized linear mixed model. The likelihood structure now accounts for PCR effects, while remaining 3C measurement effects and spatial patterns are captured in the structural terms of the model. This approach allows us to more confidently call significant interactions from 4C-seq experiments, and identify differential interactions between 4C-seq experiments.

The significant interactions we obtained are well-aligned with biologically relevant signals, such as CTCF occupancy and H3K27ac deposition, and recover known functional elements at key lineage-determining transcription factors. Furthermore, our estimates of differential contact frequency show high concordance with deduplicated Hi-C data. This agreement indicates that the transition to a binarized framework does not result in a substantial loss of biological information; rather, it provides a more robust estimate of the underlying 3D signal than count-based methods. We then examined the relationship between enhancer function and looping by targeting the functional enhancer(s) of *GATA6* and *ONECUT1* with CRISPR. Our results indicate that looping to the promoter can occur during early differentiation stages even with a non-functional enhancer, but at the moment of gene induction, the contacts fail to accumulate in the presence of a non-functional enhancer.

Nevertheless, our method assumes that there is sufficient spatial variation in the zeros to permit estimation. In regions where capture probabilities are consistently high, our estimator will be unable to resolve differences in capture rates because it only sees the presence/absence indicator. Next, if capture rates are large, such that capture probabilities are close to one, a zero is no longer sufficiently informative, and the complementary log-log link function returns an infinite estimate. This may occur if large amounts of genomic DNA are initially used in the 4C-seq reaction. Finally, despite the speed of INLA, this method is restricted to local windows around the bait, but we have successfully run this method on regions up to 5 Mbp.

## Conclusions

By reconceptualizing 4C-seq analysis at the fragment level under binarization, FourC simultaneously preserves the high-resolution nature of the assay and addresses the assay’s primary technical limitation: PCR duplication of counts. While binarization eliminates a technical artifact of the measurement process, it weakens the informa-tiveness of individual fragment-level measurements. To overcome this, we utilize a Gaussian process prior to share information between fragments and model both coarse-and fine-grained spatial patterns to recover the underlying biological signal.

We demonstrate that FourC enables us to better estimate chromatin contact frequencies across many 4C-seq experiments. This provides two key advantages: first, it allows us to better prioritize functional genomic elements compared to existing methods, and second, it returns more accurate differential interactions when compared to Hi-C data. By focusing our estimation efforts on a signal devoid of PCR inflation, we find that our method more efficiently detects differences due to CRISPRi, which were previously obscured by count-based methods, and is sensitive to small local contact differences.

By obtaining estimates of contact frequency without PCR artifacts, 4C-seq can be better analyzed and integrated with other data modalities to further study the interplay between gene expression, enhancer function, and chromatin topology. Beyond 4C-seq, the model remains readily applicable to other 3C data that admit a 1D representation by replacing the Bernoulli likelihood with a Poisson or negative binomial likelihood to identify significant interactions and differential interactions.

## Methods

### hPSC culture and directed differentiation

Experiments were performed with H1 human embryonic stem cells (NIHhESC-10-0043), which were regularly confirmed to be mycoplasma-free by the Memorial Sloan Kettering Cancer Center (MSKCC) Antibody & Bioresource Core Facility. All experiments were conducted per NIH guidelines and approved by the Tri-SCI Embryonic Stem Cell Research Oversight (ESCRO) Committee. hPSCs were maintained in Essential 8 (E8) medium (Thermo Fisher Scientific, A1517001) on vitronectin (Thermo Fisher Scientific, A14700) pre-coated plates at 37°C with 5% CO2. The Rho-associated protein kinase (ROCK) inhibitor Y-276325 (5 µM; Selleck Chemicals, S1049) was added to the E8 medium immediately after passaging or thawing of hPSCs for 24 hours.

hPSCs were initially seeded at ∼0.75-1E6 cells/well in a 6-well plate and maintained in E8 medium for 2 days to reach ∼80% confluence. Cells were washed with PBS and differentiated to DE, FG, and PP following previously described protocols [24]. hPSCs were rinsed with PBS and first differentiated into DE using S1/2 medium supplemented with 100 ng/ml Activin A (Bon Opus Biosciences) for 3 days and CHIR99021 (Stemgent, 04-0004-10) for 2 days (first day, 5 µM; second day, 0.5 µM). DE cells were rinsed with PBS and then exposed to S1/2 medium supplemented with 50 ng/ml KGF (FGF7) (PeproTech, 100-19) and 0.25 mM vitamin C (VitC) (Sigma-Aldrich, A4544) for 2 days to reach FG stage. FG cells were then switched to S3/4 medium supplemented with 50 ng/ml FGF7, 0.25 mM VitC and 1 µM retinoic acid (RA) (Sigma-Aldrich, R2625) for 2 days to reach PP stage.

### 4C-seq

For each primary digestion reaction, approximately 10 × 10^6^ cells were fixed with 10 ml of 1% formaldehyde in PBS for 10 min at room temperature. The reaction was quenched with ice-cold 1M glycine, and cells were collected by centrifugation (500 ×*g*, 5 min, 4^◦^C). Pellets were washed with PBS and stored at −80^◦^C. Cells were resuspended in ice-cold lysis buffer (10 mM Tris pH 7.5, 10 mM NaCl, 0.2% NP-40, with protease inhibitors) and incubated for 15 min on ice. Following centrifugation, the pellet was washed with 1.2x digestion buffer and resuspended in 1.2x buffer containing 0.3% SDS, then incubated for 1 hr at 37^◦^C while shaking at 750 rpm. SDS was quenched by adding 20% Triton X-100 and incubating for 1 hr at 37^◦^C.

The samples were treated with the primary restriction enzyme and incubated overnight at 37^◦^C. Digestion efficiency was verified by electrophoresis, detecting a smear between 0.2 and 2 kb in a 1.5% agarose gel. The enzyme was deactivated (65^◦^C, 20 min, 750 rpm), and ligation of DNA ends was performed in T4 ligation buffer with T4 DNA Ligase overnight at 16^◦^C. Ligated samples were treated with Proteinase K and reverse crosslinked overnight at 65^◦^C. Following RNase treatment, phenol/chloroform extraction, and DNA precipitation, pellets were dissolved in 10mM Tris pH 8.

For the second digestion, a secondary enzyme was added to the template and incubated overnight at 37^◦^C. The enzyme was inactivated at 65^◦^C for 20 min, and a second ligation was performed overnight at 16^◦^C. The final ligation product was extracted by phenol/chloroform and quantified via fluorometric assay. For library preparation, viewpoint-specific primers were designed around the promoter of interest. Library preparation was performed using 200 ng of 4C-template DNA via PCR (94^◦^C for 2 min; 16 cycles: 94^◦^C for 10 s, 59^◦^C for 1 min, 68^◦^C for 3 min; final extension: 68^◦^C for 5 min).

Amplified DNA was cleaned using magnetic beads. A second round of PCR was performed using the amplified DNA as a template to add sequencing primers and indexes (94^◦^C for 2 min; 20 cycles: 94^◦^C for 10 s, 60^◦^C for 1 min, 68^◦^C for 3 min; final extension: 68^◦^C for 5 min). 4C PCR library efficiencies were confirmed by automated electrophoresis and sequenced in PE100/PE150 mode.

### Hi-C

Two million cells were collected and fixed with 1% formaldehyde. The subsequent steps of Hi-C were then performed using the Arima-Hi-C kit (Arima, A510008), while libraries for sequencing were prepared with the KAPA Hyper Prep Kit (KAPA, KK8502) following the manufacturers’ guidelines. Samples were pooled and submitted to MSKCC Integrated Genomics Operation core for quality control and sequencing on Illumina HiSeq 4000 platform.

### CRISPR

CRISPRi experiments were performed as previously described against *GATA6* +e9 and +e12 [13]. CRISPR-Cas9 experiments were performed as previously described against *ONECUT1* +e664 [14]. Refer to their respective papers for details on guide design and generation of CRISPR lines.

### 4C data processing and analysis

Raw FASTQ files were supplied to pipe4C with the relevant restriction enzymes used for the primary and secondary digest steps [3]. Different experiments utilized different combinations of restriction enzymes (NlaIII, DpnII, MboI, Csp6I). Reads were aligned against hg38, with all alternative haplotypes removed with Bowtie 2.4.1. All blind and non-blind fragments were reported over the entirety of the genome. Otherwise, default settings were used to process the fastqs. Only fragment-level count summaries were used, and no normalization was used.

peakC 1.02 was used to identify significantly interacting regions at an FDR of 0.1 with a window size of 21. FourSig 2.2.0 was similarly applied at an FDR of 0.1 utilizing a default window size of 1. Because pipe4C returns pre-formatted files containing all valid fragments, FourSig was run directly on the position and read count data. Lastly, r3Cseq 1.8.0 was used to call significant interacting fragments under default settings; to facilitate this, its core significance-calling function was executed directly on the pipe4C output rather than through the standard function wrappers.

For testing for differential regions between conditions with DESeq2, the window of interest around each viewpoint was tiled into 25 kb bins. Next, the total counts corresponding to the fragments that fall into each bin were summed to construct the count matrix per replicate. DESeq2 1.40.1 was then run on these count matrices to identify differential contacts between conditions using the 4C reads.

### Hi-C data processing and analysis

Prior to alignment, Hi-C Pro 2.11.4 was used with GATC and GANTC as the restriction sites, with all alternative haplotypes removed. Reads were aligned with default settings for Bowtie 2.4.1 in Hi-C Pro, for both the global and local alignment steps. After both alignment steps, reads with a MAPQ of at least 30 were retained for further analysis and duplicates were removed. Sample-level Hi-C maps were converted to .Hi-C file format with JuicerTools 1.22.01. For each subregion presented in the experiments above, we extracted the counts at 25 kb from the underlying Hi-C files with strawr 0.0.91 and then used DESeq2 1.40.1 to identify differential contacts between conditions at an FDR of 0.1.

### ChIP-seq data processing and analysis

Processed H3K27ac ChIP–seq signal tracks aligned to hg38 (bigWig format) were obtained from GEO accession GSE114102 (Huangfu lab). CTCF ChIP–seq bigWig tracks aligned to hg38 were downloaded from GEO accession GSE211101 as deposited by the original study. Tracks were visualized in R; no additional processing was applied.

### Binary transformation and estimation

Under the model that *o_i_* ∼ *Ber*(1 − *e*^−^*^λi^* ), each fragment will occur with probability 1 − *e*^−^*^λi^* . Under the hypothesis that *λ* is approximately homogeneous in some small window, it becomes possible to furnish an estimator for the average capture rate 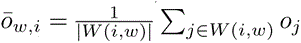. Then under approximate homogeneity, 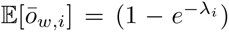 Furthermore, the variance of the estimator is given as 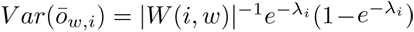. Crucially, the properties of this estimator only depend on the underlying capture rate at the original step of the 4C-seq protocol and do not depend on PCR effects.

Finally, observe that this quantity is a monotonic transform of the capture rate function, *λ*, and hence it will preserve relative relationships between observations. This suggests that merely assessing the overall capture rates inside a smoothed running window is a way to view the data without duplication effects, though technical effects owing to the underlying library are still present, such as the number of cells being used, but vanish under similar experimental parameters.

### Specification and estimation of spatial generalized linear mixed model

For a given collection of experiments collected from the same bait, let 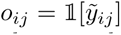 be the binary capture indicator where *i* is the *i*th fragment in the region under consideration, and *j* is the *j*th replicated measurement. For each fragment, we have *x_ij_* a collection of covariates pertaining to the measurement *o_ij_*. Finally, the data can be spatially indexed by *s_ij_*, which is the position of the *i*th fragment. Our data thus consists of (*o_ij_, x_ij_, s_ij_*), for *i* ∈ [1*, …, F*]*, j* ∈ [1*, …, J*], where we have a total of *F* fragments, up to say 1.2 Mb, around the bait and *J* replicates in the experiment.

Fixed effects are used to model technical effects, such as fragment-level characteristics, such as GC, length, type, and sample origin. GC and length covariates are to be modeled using spline terms to yield natural splines with 3 degrees of freedom. All terms are assigned flat Gaussian priors (low precision). The remaining (spatial) random effects of our model are assigned a Matern Gaussian Process prior. The behavior of the Matern process is controlled by *σ*^2^*, ρ, ν*, which correspond to the marginal standard deviation, the range, and the smoothness respectively. *ν* is typically not identified by data and will be fixed at *ν* = 2, leaving two free parameters during estimation [25]. These parameters are assigned a penalized complexity prior to shrink our Matern Gaussian process towards a constant function [26]. Finally, the realization of the Matern process is allowed to vary across different experimental conditions.

Although one spatial term could be sufficient, if there are multiple scales of contacts that exist, it would be better to model both short- and long-range contact patterns. For example, the overall characteristic distance decay is removed because it is not thought to yield much useful information regarding the important regulatory contacts [20, 21]. To that end, we leverage ideas from Lattice Kriging (LK), and for data arising from the exponential family, Extended Lattice Kriging (ELK) [27, 28]. Essentially, the 2D domain of interest is partitioned into a fine and coarse lattice, and then continuous simultaneous autoregressive priors with short and long correlation length scales are placed on the terms in the lattice. This captures both long- and short-range dependencies within the data and is shown to be more adaptive for capturing spatial heterogeneity.

A similar idea may thus be leveraged here in the case of spatially smooth terms that have been assigned a Matern prior, given that the Matern process will be ultimately approximated via a basis expansion. To do this in the 1D setting, the 1D domain is divided into an evenly spaced grid. Adopting the terminology from splines, the interior knots are placed uniformly along the domain, and compactly supported triangular basis functions are placed at each knot. The first knotting sequence will be realized at a coarse resolution, *δ^Coarse^* = 50 kb, and the second knotting sequence at a finer resolution of *δ^F ine^* = 10 kb. This proposed construction unites previous modeling efforts that condition on long-range patterns but simultaneously models short-range patterns. For example, in the context of Hi-C, the observed-over-expected matrix removes the large-scale distance decay [29].

#### Observation model

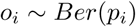

#### Latent Gaussian model

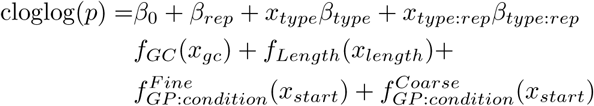

#### Priors/hyperpriors

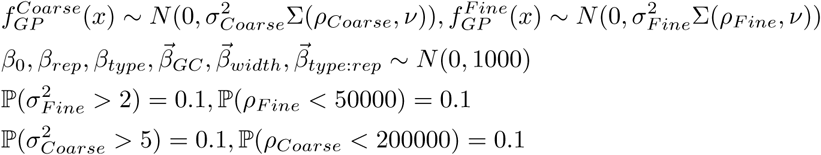

As our model admits a formulation as a latent Gaussian model (LGM), we may use the Integrated Nested Laplace Approximation (INLA) to approximate the posterior distribution of all model quantities. In comparison to alternative Gaussian process kernels, the Matern Gaussian kernel is a solution to a stochastic partial differential equation (SPDE) and can be used to derive a sparse precision matrix. These advances are packaged into *inlabru* [30–33] and make estimation quick and straightforward. We use inlabru 2.12.0 for model estimation.

### Posterior processing of spatial generalized linear mixed model

In the INLA scheme, the approximated posterior density is obtained by using a discrete grid over the sets of hyperparameters, each one having weights proportional to the marginal likelihood under the setting at the specified node. Let *w_i_* be the weight of the *i*th hyper-parameter setting, to obtain draws from the full posterior density, one first draws *θ* ∼ *Cat*(*w_i_*), and subsequently, a draw from *x* ∼ *p*(*x*|*y, θ*) is obtained to yield a final posterior draw. This operation is repeated *B* times to yield a matrix of posterior draws for the parameters of interest in the form of a *N* × *B* matrix for the *N* parameters of interest. These parameters can then be subjected to arbitrary transformations, and inference can proceed on the quantities of interest, such as the difference between parameters in the model, e.g. differences in spatial trends between conditions. The *inlabru* package provides an interface for generating samples from each of the elements of the nodes of the computation graph.

#### Gaussian excursions

The problems of identifying significant interactions and differential contacts are connected in the sense that in the former, we are seeking an enrichment above a chosen background, and in the latter, we are seeking an enrichment or depletion with respect to another experiment. Both of these ideas can be conceptually joined through the ideas of Gaussian excursions [23]. Abstractly, we consider the spatial function, *f*(*s*) that acts on a spatial domain and reveals the intensity of a signal. In typical settings, we may be interested in evaluating the degree to which ℙ(*f*(*s_i_*) ≥ *u*|*y*) holds, and in quantifying our uncertainty in this belief. This can be carried out position by position, but if joint significance is required, we will need to discuss *excursion sets*.

For *f*(*s*) : Ω → ℝ, a mapping from the spatial domain of interest to the real numbers, let 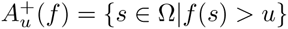, and analogously for 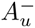. An excursion set is defined for a random process *x*(*s*) as 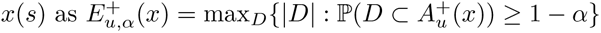. This gives the largest set of indices across Ω within *D* that are in the excursion set 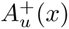 with at least probability 1 − *α* [23].

This defines the set of indices where all values of the random process simultaneously exceed the chosen reference value. In the case of significant and differential interactions, this reference value is chosen to be 0, as explained below. The joint distribution of the posterior draws is thus used to assess the probability that all chosen nodes simultaneously exceed u, and its computations can be performed through the *excursions* package [34]. Both marginal and joint tests are available in our package.

#### Enrichment analysis for significant interactions

A significantly interacting region is typically defined with respect to some fixed background. The background can either be locally or globally defined [20, 21]. Under the decomposition of the spatially smooth trend into two components, one coarse and the other fine, the fine-scale term can be thought of as the small-scale fluctuations that exist after removing the large-scale trend. The prior placed on the fine-scale component encourages similarity of neighbors. When this fine-scale term is 0, it reverts to the large-scale trend, but when it is not, it is enriched above the large-scale local trend. In other words, f*_F ine_*(s*_i_*) yields a very convenient way to define a significant interaction: check if p(f*_F ine_*(s*_i_*) > 0|y).

#### Fold changes for differential interactions

Similarly, a differentially interacting region can be defined by comparing the spatially smooth trends Δ(*s_i_, A, B*) = (*f_B,Fine_*(*s_i_*) + *f_B,Coarse_*(*s_i_*)) − (*f_A,F ine_*(*s_i_*) + *f_A,Coarse_*(*s_i_*)). To make clear how this refers to a fold change, consider a piecewise constant intensity for the overall biological rate. Then in a standard log-linear model 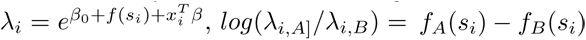, since all other things being equal, the only value that changes is the spatially varying signal. Thus the test becomes identical: *p*(Δ(*s_i_, A, B*) *>* 0|*y*). To perform a two-tailed test, we check if the 1 − *α* credible interval contains 0. We will now argue that 0 is a meaningful reference level. Spatial terms are constrained so that Σ*_i_ f*(*s_i_*) = 0 for identifiability reasons. Hence, by definition Σ*_i_ f_A_*(*s_i_*)−*f_B_*(*s_i_*) = 0, which implicitly requires that the average fold change is 0. This constraint mimics the constraint of standard differential count methods, which assume that robust measures of the average fold change is 0 [35, 36]. Thus, it will be anticipated that using this implicit normalization, the estimates will agree with existing count methods, and partially explains the concordance of the Hi-C and 4C estimates for fold change estimation.

## Declarations

### Availability of data and materials

Datasets generated in this study will be deposited in GEO prior to publication. Accession numbers will be provided upon availability. Publicly available datasets used in this study are GSE114102 and GSE211101. The FourC software package is available at FourC.

### Authors’ contributions

WW conceived and developed the statistical methodology, implemented the software package, processed the data, and performed the analysis. SJK and JY designed and performed *ONECUT1* experiments. RL designed and performed *GATA6* experiments. JP designed and performed *PDX1* experiments. CSL and DH supervised the study. WW and CSL wrote the manuscript with input from all authors.

## Acknowledgements

We thank Rui Yang, Feiyang Huang, Effie Apostolou, Wesley Tansey, and Kushal Dey for valuable discussions and feedback that helped refine the statistical methodology. We would also like to thank Erin Char and Ukjin Lee for testing the package.

This work was supported in part by National Institutes of Health grants U01HG012051 (DH), U01DK128852 (CSL and DH), and T32GM132083 (WW).

