## Supplemental Information for "FourC: identifying significant and differential contacts in 1D chromatin conformation data"

### Supplementary Information for: FourC: identifying significant and differential contacts in 1D chromatin conformation data

Wilfred Wong[1,2], Samuel J Kaplan[3], Renhe Luo[3], Julian Pulecio[3], Jielin Yan[3], Danwei Huangfu[3], Christina S. Leslie[1*]

[1]Computational and Systems Biology Program, Memorial Sloan Kettering Cancer Center, 417 E 68th St, New York, 10065.

[2]Tri-Institutional Program in Computational Biology and Medicine, Weill Cornell Medical College.

[3]Developmental Biology Program, Memorial Sloan Kettering Cancer Center, 417 E 68th St, New York, 10065.

Contributing authors:;

**Abstract**

This file contains the Supplementary Note (statistical derivations for the binarization and PCR-duplication model), Supplementary Figures S1–S17, and Supplementary References for the manuscript "FourC: identifying significant and differential contacts in 1D chromatin conformation data."

### Supplementary Note

#### Chromatin contacts, binarization, and regression

##### Multinomial analysis of chromatin capture frequencies

3C experiments assay the spatial proximity of two genomic loci by measuring the occurrence of proximity ligation events. The products of a 4C-seq reaction can be modeled as $y \sim \text{Multi}(N_c, p_G)$ since over $N_c$ cells, the contacts exhibit relative preferences according to $p_G$. To encode spatial effects, $s_k$, and technical effects, $x_k$, we model relative preferences as $p_{G,k} = e^{\eta_k} / \sum_j e^{\eta_j}$ where $\eta_k = s_k + x_k^T \beta$ and $s_k$ is locally smooth. Under the multinomial assumption, an equivalent Poisson model simplifies analysis [1–3]. Defining $y_k \sim \text{Pois}(e^{\eta_k})$, and $N = \sum_k y_k \sim \text{Pois}(\tau), \tau = \sum_j e^{\eta_j}$, the joint Poisson likelihood, $L_y$, is equal to the product of a multinomial, $L_{y|N}$, and Poisson likelihood, $L_N$ (Eqs. 1)

$$\begin{aligned} L_{y|N}(\eta) L_N(\tau) &= \left[ \binom{N}{y_1, \ldots, y_K} \prod_i \left( \frac{e^{\eta_i}}{\sum_j e^{\eta_j}} \right)^{y_i} \right] \frac{e^{-\tau} \tau^N}{N!} \\ &= \prod_i \frac{(e^{\eta_i})^{y_i} e^{-e^{\eta_i}}}{y_i!} = L_y(\eta) \end{aligned} \quad (1)$$

Now, we give some arguments for why $s_k$ is the object of interest in 3C assays. First, we extend our previous model to 2D indices, as in Hi-C, and collapse fragments into bins, so that $i, j$ index genomic bins instead, $s$ plays the anticipated role, and bias effects are symmetric in the index $y_{ij} \sim \text{Pois}(e^{\beta_0 + s_{ij} + x_i^T \beta + x_j^T \beta})$. To remove common bias effects along indices $i, j$, Iterative Correction and Eigenvector decomposition (ICE) normalization estimates the bias associated with each linear genomic index to obtain $T_{ij} \propto e^{s_{ij}}$. A similar strategy is present in the vanilla-coverage normalization method [4, 5].

Next, we assess the computation of fragment-level fold changes in 4C. A log-linear model for the $i$th fragment is given as $y_{ij} \sim \text{Pois}(\exp(\beta_0 + s_{ij} + x_i^T \beta))$, where $j$ indicates the experimental condition of the observation (not the second spatial index as in Hi-C). Because the fragment-specific technical covariates, $x_i$, are strictly constant across conditions, the log fold change between conditions depends only on the spatial parameter: $\log(\lambda_{ij} / \lambda_{ij'}) = s_{ij} - s_{ij'}$. Finally, in both 4C and Hi-C, aggregation proceeds logically. If the spatial signal $s$ is nearly constant locally, the expected sum of counts for two neighboring fragments, $i$ and $i'$, averages over technical effects: $e^{\beta_0 + s}(e^{x_i^T \beta} + e^{x_{i'}^T \beta})$.

The analysis above applies to the latent molecular products $y_i \sim \text{Pois}(\lambda_i)$, which are never directly observed. $y_i$ is subjected to library preparation steps, such as PCR, prior to sequencing to generate the observed read counts, $r_i$, per experiment. Under these assumptions, we now examine a minimal model that captures how those library preparation effects propagate into $r_i$ and why binarization should be applied to $r_i$. To build intuition, we first examine the simpler problem of positive duplication acting on a Poisson random variable.

#### Bernoulli and Poisson random variables

Suppose that $y_i \sim \text{Pois}(\lambda_i)$ for $i \in [1, ..., n]$ but we only observe $\tilde{y}_i = \nu_i y_i$, with $\nu_i \in \mathbb{Z}^+$, which is the original data subject to some arbitrary positive duplication. Assume $\nu_i$ is fixed and define the binary random variable $z_i = \mathbb{1}[\tilde{y}_i = 0]$. Since $\nu_i > 0$, $z_i = \mathbb{1}[\tilde{y}_i = 0] \equiv \mathbb{1}[y_i = 0]$. Thus, to estimate $\log(\lambda_i)$, we can use the estimator $\widehat{\log(\lambda_i)} = \log(-\log(\hat{p}))$, where $\hat{p}$ is the observed proportion of zeros (Eq. 2).

A few properties of this estimator are as follows: i) it will consistently estimate $\lambda_i$; ii) it has a variance larger than that of the standard Poisson; iii) it fails to exist if $\hat{p} = 0$, which occurs when $\lambda_i$ is sufficiently large; and iv) it does not depend on the unknown $\nu_i$. In practice, because 4C-seq experiments typically contain only up to two replicates, stable estimation of $\hat{p}$ requires pooling information across replicates or spatially adjacent measurements [6].

$$z_i = \begin{cases} 1 & \tilde{y}_i = 0 \\ 0 & \tilde{y}_i > 0 \end{cases} \equiv \begin{cases} 1 & y_i = 0 \\ 0 & y_i > 0 \end{cases} \rightarrow z \sim Ber(e^{-\lambda_i}) \tag{2}$$

#### Complementary log-log binomial and log-linked Poisson regression

Now, suppose $y_i$ are related to fixed $x_i$ by $y_i|x_i \sim \text{Pois}(e^{x_i^T \beta})$. $\beta$ is estimated through a log-linked Poisson generalized linear model (GLM), where $\log(\lambda_i) = x_i^T \beta$. Under positive count duplication $\tilde{y}_i = \nu_i y_i$, we define the presence/absence indicator $o_i = \mathbb{1}[\tilde{y}_i > 0]$, so that $o_i \sim \text{Ber}(p_i)$. Consequently, $\nu_i$ can be eliminated by the complementary log-log link function, where $\widehat{\log(\lambda)} = \log(-\log(1 - \hat{p}))$ (Equation 3). The cloglog link is not the canonical link function, but has previously been used to assess Poisson rates [7, 8].

$$\begin{aligned} \text{cloglog}(p_i) &= \log(-\log(1 - p_i)) = \log(-\log(1 - 1 + e^{-\lambda_i})) \\ &= \log(-\log(e^{-\lambda_i})) = \log(\lambda_i) = x_i^T \beta \end{aligned} \tag{3}$$

Finally, we note that the cloglog link is "natural" in the analysis of discrete Poisson point processes. Consider a 1D inhomogeneous Poisson point process (IHPP). In that setting, the number of points falling in an interval is given by $y_i \sim \text{Pois}(\int_{D_i} \lambda(s) ds)$, where $\lambda$ depends on $x$. If only presence/absence data are available, estimating the parameters of the IHPP is best done with a cloglog-linked binomial regression [6].

In this section, we have established a model for the molecular products of a 4C-seq reaction, and shown that it can be approximated by a Poisson distribution. Subsequently, $s$, the spatial parameter, $s_i$, is the parameter of interest, and naturally arises in existing estimation and inference schemes. We then proved that under arbitrary positive count duplication, presence/absence indicators naturally recover the underlying Poisson rate $\lambda_i$ via the complementary log-log relation, effectively eliminating PCR duplication artifacts.

#### High throughput sequencing statistical modeling

Having now shown the link between $y_i$ and $o_i$, it is clear that for any duplicated data, estimation and inference are straightforward. Nevertheless, the remaining difficulty is that what we observe is not $\tilde{y}_i$, but $r_i$, the result of preparing a library (4C-seq), duplicating (PCR), and sequencing. To understand the degree to which our previous analysis holds, we will model how $y_i$ is transformed into $r_i$. PCR is controlled by the efficiency of the reaction, $p_{d,i}$, and the number of cycles, $c$, so that the final number of counts arising from $y_i$ is given as $\rho_i = \text{PCR}(y_i, p_{d,i}, c)$. Subsequently, sequencing selects molecules without replacement from the prepared library $L = \{\rho_i\}$ to obtain $S = \{r_i\}$. If $\mathbb{P}(r_i = 0) \approx e^{-\lambda_i}$, $\mathbb{P}(o_i = 0) = e^{-\lambda_i}$ will also hold for binarized read counts.

#### Duplicating molecular products as a Galton-Watson process

First, instead of assuming that molecules are duplicated according to a fixed $\nu_i$, we model PCR as a stochastic branching process. In a single amplification cycle, each molecule is successfully copied with probability $p_{d,i}$. If it fails to replicate, only the original copy is retained. Therefore, the number of molecules after one cycle given an initial count $y_i$ is $y_i + \delta_i$, where $\delta_i \sim \text{Bin}(y_i, p_{d,i})$. Iterating this process for $c$ cycles over the products of the prior cycle, and assuming no losses, the amplification strictly obeys a Galton-Watson (GW) branching process [9]. The final amplified molecule count, $\tilde{y}_i$, is thus distributed as $\text{GW}(y_i, p_{d,i}, c)$ (Eq. 4).

$$\text{PCR}(y_i)|y_i = (y_i + \sum_{j=1}^{y_i} d_i) \rightarrow \tilde{y}_i(c, p_{d,i}) = \underbrace{\text{PCR}(\text{PCR}(...\text{PCR}(y_i)))}_{c} \tag{4}$$

Crucially, because the branching process cannot generate a count if the molecule was never present, $y_i = 0 \Rightarrow \tilde{y}_i = 0$. This fact depends only on lossless copying, not on the specific branching process. Other PCR models could be adopted that account for synthesis errors and size-dependent branching processes [10, 11]. Regardless, the model outlined here captures the essential characteristics of PCR counts: i) PCR is a stochastic process, and ii) PCR preserves zeros.

Finally, if we seek to estimate $\lambda_i$, we could attempt to marginalize over the randomness of the GW process, but this is analytically difficult and computationally intractable. Instead, observe that factorizing the marginal likelihood of $\tilde{y}_i$ yields a simplification that bypasses the nuisance parameters $p_{d,i}, c$ (Eq. 5) [12]. In other words, the presence/absence indicator retains sufficient information to estimate $\lambda_i$, while the discarded information is confounded by PCR effects:

$$\tilde{y}_i|y_i \sim \begin{cases} 0 & y_i = 0 \\ \text{GW}(y_i, p_{d,i}, c) & y_i > 0 \end{cases} \rightarrow p(\tilde{y}_i|\lambda_i, p_{d,i}, c) = (e^{-\lambda_i})^{\mathbb{1}[\tilde{y}_i=0]} p(\tilde{y}_+|\lambda_i, p_{d,i}, c)^{1-\mathbb{1}[\tilde{y}_i=0]} \tag{5}$$

#### Sequencing molecular products as a multivariate hypergeometric distribution

Library preparation consists of generating $\{y_i\}$ from a population of cells and then amplifying the genomic material to yield the library $L = \{\tilde{y}_i\}$. Subsequently, the sequenced reads, $S = \{r_i\}$, are obtained by drawing $N_R$ reads from $L$ without replacement according to the relative concentrations of the starting molecules. This step is formally described by the multivariate hypergeometric (MVHG) distribution [2, 13].

In this section, we temporarily assume a fixed duplication constant, $\nu_i$, before incorporating full PCR effects. Because the total number of sampled reads represents only a small fraction of the amplified library ($N_R \ll |L|$), we can accurately approximate the MVHG with a multinomial distribution, $\mathrm{Multi}(N_R, \gamma_i)$, where $\gamma_i = \tilde{y}_i/\tilde{y}_+$. Crucially, this sampling process ensures that if the initial count $y_i = 0$, it is maintained as a zero.

The multinomial correlations will often lead to difficult computations, and so we further approximate the final read counts by a Poisson $r_i|\tilde{y}_i, \nu_i \sim \mathrm{Pois}(\frac{N_R}{\tilde{y}_+}\nu_i y_i)$. As the genome is quite large, the individual capture probabilities will tend to be small, and so, the dependence between $y_i$ and the total library sum $\tilde{y}_+$ is negligible. Therefore, treating $\tilde{y}_+$ as a constant with respect to $y_i$ yields approximately independent observations.

Following that approximation, Equation 6 indicates that provided $\nu_i$ is sufficiently large, and $R = N_R/\tilde{y}_{+,i}$ is not too small, the major contributor to the zero probability will arise from the fact that the original value had a strong tendency towards zero. For example, standard protocols suggest 46 cycles of PCR, 5 million cells, and 1 million reads [14]. Assuming $R \approx 10^{-10}$ and perfect PCR efficiency ($\nu_i = 2^{46}$), the exponent term $e^{-\nu_i R} \approx e^{-7000} \approx 0$. Under these conditions, detecting a molecular species even once in the sequenced library provides overwhelming evidence of its presence. A simulation demonstrating this property is presented in Fig. S1, which shows the agreement between simulation and theory.

$$\begin{aligned}
\mathbb{P}(r_i = 0|\nu_i, N, \tilde{y}_+) &= \sum_j \mathbb{P}(r_i = 0|\nu_i, N, \tilde{y}_+, y_i = j)\mathbb{P}(y_i = j) \\
&= \sum_{j=0}^{\infty} e^{-\nu_i R j} \frac{e^{-\lambda_i}\lambda_i^j}{j!} \\
&= e^{-\lambda_i} \sum_{j=0}^{\infty} \frac{(e^{-\nu_i R}\lambda_i)^j}{j!} \\
&= e^{-\lambda_i(1-e^{-\nu_i R})}
\end{aligned} \tag{6}$$

Hence, our arguments in the previous section continue to hold. Under fixed positive count duplication and subsampling from a prepared library, binarization continues to allow us to estimate $\lambda_i$ even if $\nu_i$ is unobserved. We now turn our attention to random duplication by a Galton-Watson process.

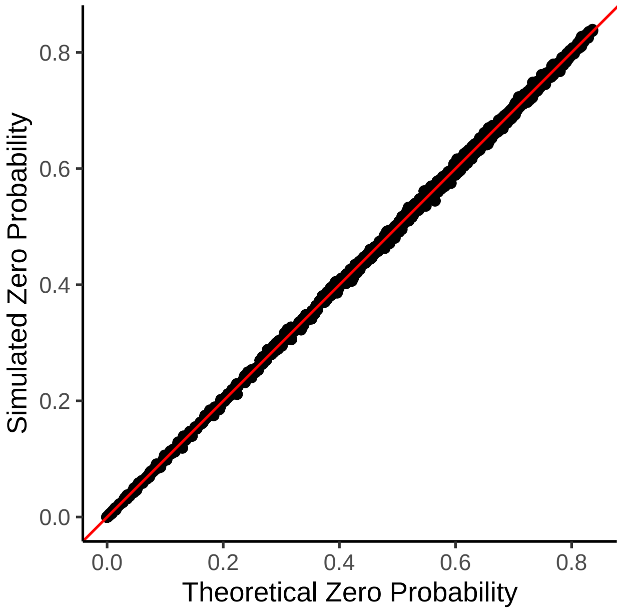


**Fig. S1**: Simulation of generating 0 counts vs. theoretical zero probability.

#### Two-stage modeling for sequencing duplicated molecular species

In the prior two sections, we outlined the implications of duplication and sequencing on the detection of a zero under two parallel regimes. First, we established that a stochastic branching process strictly preserves biological zeros during PCR amplification. In the subsequent section, we demonstrated that under fixed but sufficient duplication, the probability of observing a zero after library subsampling is almost entirely driven by the initial absence of the target molecule ($y_i = 0$). We now unify these two parallel regimes and demonstrate that under both random amplification and sequencing, $\mathbb{P}(r_i = 0) \approx e^{-\lambda_i}$. Consequently, for the presence/absence indicator $o_i = \mathbb{1}[r_i > 0]$, the approximation $\mathbb{P}(o_i = 1) \approx 1 - e^{-\lambda_i}$ holds, so that the complementary log-log link can be used to estimate $\lambda_i$.

We are now ready to state the full ideal model: i) $y \sim \text{Multi}(N_c, p_G)$ models the initial read counts in the cell population; ii) $\rho_i | y_i \sim \text{GW}(y_i, p_{d,i}, c)$ duplicates the molecules of each $y_i$ independently of the other observations, but returns 0 if $y_i = 0$; and iii) $r_i | \rho_i \sim \text{MVHG}(\rho_i, N_R)$ models sampling from the sequencer with $N$ reads. From the point of view of estimation, there are multiple problems: i) the multinomial induces correlations across $y_i$; ii) the Galton-Watson process has a difficult-to-utilize probability mass function; iii) the multivariate hypergeometric distribution is difficult to analyze.

This leads to a series of approximations: i) replace the multinomial in the first step with $y_i \sim \text{Pois}(\lambda_i)$; and ii) replace the MVHG with a multinomial and then a Poisson approximation so that $r_i | \rho_i, y_i > 0 \sim \text{Pois}(R\rho_i(y_i, p_{d,i}, c))$. Equation 7 thus reveals that there are two places where a zero may arise: either from the preparation of the sample itself, that is $y_i = 0$, or during sampling, if we have insufficiently sampled, so that even though $y_i > 0$, $r_i = 0$. The PCR process, assuming losslessness, can only preserve existing zeros. We will now attempt to show that Equation 8 holds approximately.

$$\begin{aligned} y &\sim \text{Multi}(N_c, p_G) \\ \rho|y &\sim \begin{cases} 0 & y = 0 \\ \text{GW}(y, p^r, c) & y > 0 \end{cases} \\ r|\rho, y &\sim \text{MVHG}(\rho, N_R) \end{aligned} \rightarrow \begin{aligned} y_i &\sim \text{Pois}(\lambda_i) \\ \rho_i|y_i &\sim \begin{cases} 0 & y_i = 0 \\ \text{GW}(y_i, p_{d,i}, c) & y > 0 \end{cases} \\ r_i|\rho_i, y &\sim \begin{cases} 0 & y_i = 0 \\ \text{Pois}(\frac{N_R}{\rho_+}\rho_i) & y_i > 0 \end{cases} \end{aligned} \tag{7}$$

$$\mathbb{P}(r_i = 0) = \sum_{j=0}^{\infty} \mathbb{P}(r_i = 0|y_i = j)\mathbb{P}(y_i = j) \approx e^{-\lambda_i} \tag{8}$$

If $y_i = 0$, then $\rho(y_i, c) = 0 \rightarrow r_i = 0$. Hence $\mathbb{P}(r_i = 0|y_i = 0) = 1$ and $\mathbb{P}(y_i = 0) = e^{-\lambda_i}$. It remains to show that $\mathbb{P}(r_i = 0|y_i > 0) \approx 0$, so that zeros arise from $y_i = 0$. $\rho(y_i, c)|y_i > 0$ is described by a recursive probability distribution (Equation 9).

$$\mathbb{P}(r_i = 0|y_i = j, j > 0) = \sum_{\rho(y_i,c)=j}^{j2^c} \mathbb{P}(r_i = 0|\rho(y_i, c), y_i = j)\mathbb{P}(\rho(y_i, c)|y_i = j)$$

$$\mathbb{P}(\rho(y_i, c) = k|y_i) = \sum_{m=\lceil k/2 \rceil}^{k} \binom{m}{k-m} (p_{d,i})^{k-m}(1 - p_{d,i})^{2m-k}\mathbb{P}(\rho(y_i, c - 1) = m) \tag{9}$$

This recursive formulation is difficult to work with, so we instead consider an approximate normal distribution (Equation 10), $\rho(y_i, c)\dot{\sim}N(\mu(y_i, c), \sigma^2(y_i, c)), \mu(y_i, c) = y_i(1 + p)^c, \sigma^2(y_i, c) = y_i\frac{1-p_{d,i}}{1+p_{d,i}}[(1 + p_{d,i})^{2c} - (1 + p_{d,i})^c]$, to illustrate our point.

$$\mathbb{P}(r_i = 0|y_i = j, j > 0) = \sum_{\rho(j,c)=j}^{j2^c} e^{-R\rho(j,c)}\phi_{\mu(j,c),\sigma^2(j,c)}(\rho(j, c)) \tag{10}$$

There are now two remaining difficulties: i) $R = N_R/\rho_+$ is not independent of $\rho_i$, and ii) the summation itself. First, under a sufficient number of cycles, the individual contribution of $\rho_i$ to $\rho_+$ is small, and hence we use the same rationale as before to argue for the independence of $R$ and $\rho_i$. Next, replacing the summation with integration leads to the moment generating function (MGF) of a truncated normal, (Equation 11).

$$\mathbb{P}(r_i = 0|y_i = j, j > 0) \approx \int_{j}^{j2^c} e^{-R\rho(j,c)}\phi_{\mu(j,c),\sigma^2(j,c)}(\rho(j, c))d\rho(j, c) \tag{11}$$

This simplification yields a usable formula, given by the MGF of the truncated normal which computes the probability of observing a 0 if $y_i > 0$.

$$\begin{aligned}\mathbb{P}(r_i = 0|y_i = j, j > 0) \approx & e^{-R\mu(j,c)+R^2\sigma^2(j,c)/2} \\ & \times \left[\frac{\Phi(\beta(j,c) + R\sigma(j,c)) - \Phi(\alpha(j,c) + R\sigma(j,c))}{\Phi(\beta(j,c)) - \Phi(\alpha(j,c))}\right]\end{aligned} \tag{12}$$

To gain some intuition regarding the behavior of the reads, we consider the GW process at its mode $e^{-Ry_i(1+p)^c}$. As a function of any of its arguments, this quantity always shrinks, and exponentially tends to 0 with $c$.

As Equation 12 does not yield an easily interpretable expression, we opted to plot the behavior over various values for the parameters. As shown in Figure S2, provided that the initial counts are large, $R$ is not small, and $c = 46$, the probability that a read count is 0 is nearly 0 across various PCR efficiency regimes. This suggests that in many realistic regimes, $\mathbb{P}(r_i = 0|y_i = j, j > 0) \approx 0$.

It remains to marginalize over $y_i$ and show that the residual sum is negligible.

$$\mathbb{P}(r_i = 0) = e^{-\lambda_i} + \underbrace{\sum_{j=1}^{\infty} \mathbb{P}(r_i = 0 \mid y_i = j)\mathbb{P}(y_i = j)}_{\text{Bound}}$$

Since $\mathbb{P}(r_i = 0 \mid y_i = j)$ is observed to be monotonically decreasing in $j$, and all terms in the summation are nonnegative, we can control the sum with $\mathbb{P}(r_i = 0 \mid y_i = 1)$, which tends to 0 as $p_{d,i}$, $c$, and $R$ increase, and $\sum_{j=1}^{\infty} \mathbb{P}(y_i = j) = 1 - e^{-\lambda_i}$

$$\sum_{j=1}^{\infty} \mathbb{P}(r_i = 0|y_i = j)\mathbb{P}(y_i = j) \leq \mathbb{P}(r_i = 0|y_i = 1) \sum_{j=1}^{\infty} \mathbb{P}(y_i = j) \approx 0$$

We evaluate this marginal contribution across a range of parameter values in Figure S3 by computing a truncated sum. As expected, under standard protocols ($c = 46$ cycles), the probability of completely missing a present molecule is vanishingly small across all configurations. However, under theoretically inefficient PCR regimes, highly abundant species (large $\lambda$) exhibit the greatest sensitivity to low efficiency because their probability of a non-zero count is high. Conversely, for rare species, the underlying Poisson mass is already heavily concentrated at $y_i = 0$. Consequently, the remaining weights on the positive terms in the infinite sum are small.

To combine all of these remaining results, a Monte Carlo simulation following a multinomial, Galton-Watson, and then a MVHG sampling scheme is presented in Figure S4, where the expected number of zeros is determined almost entirely by $e^{-\lambda_i}$, the Poisson approximation to the multinomial in the first stage of sampling [2, 13].

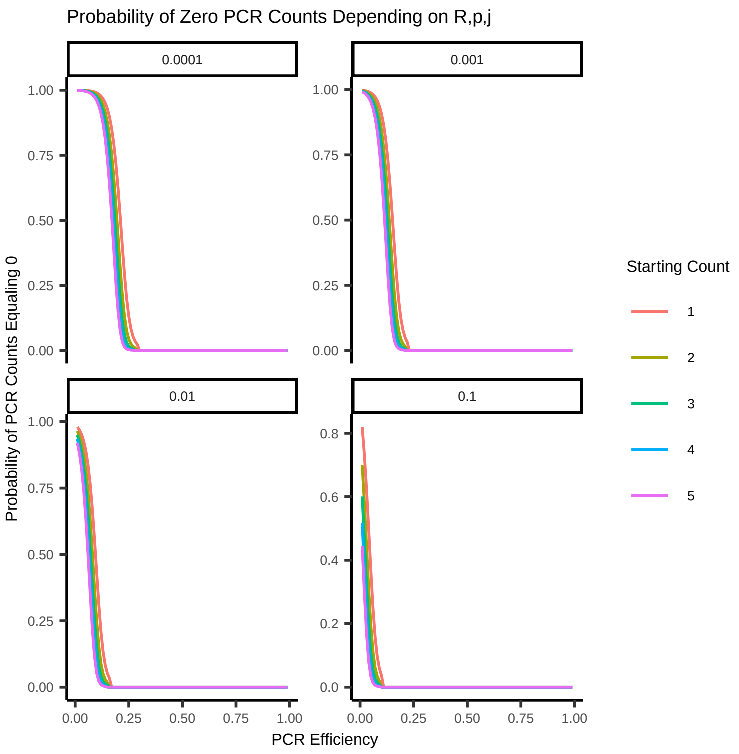


**Fig. S2**: The probability that a read count is 0 as a function of the initial starting counts. Each facet varies the total read fraction sequenced. The capture probability is plotted as a function of PCR efficiency over varying values of the initial count.

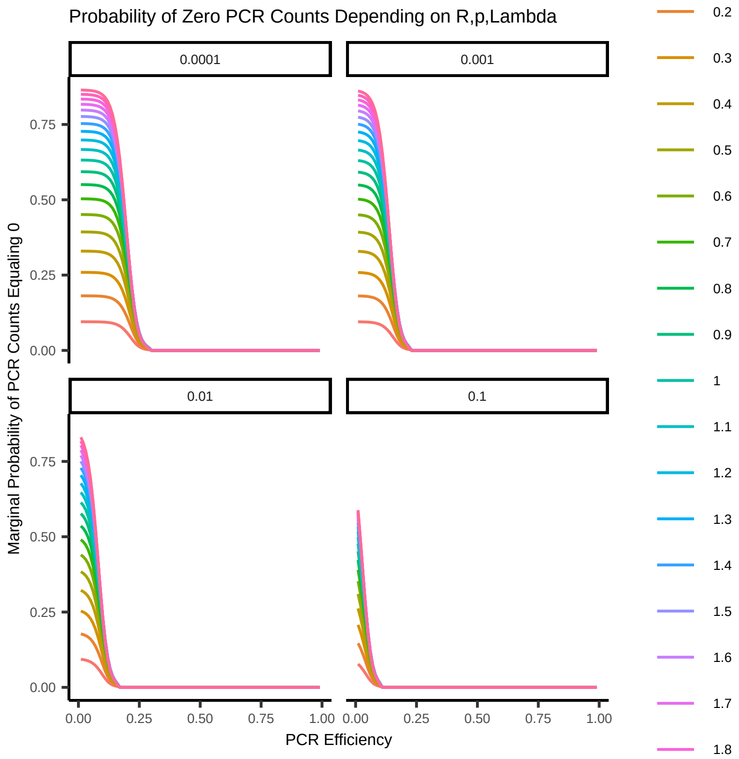


**Fig. S3**: Probability of not observing a count due to sampling effects. Each facet varies the total read fraction sequenced. The capture probability is plotted as a function of PCR efficiency over varying values of the initial rate.

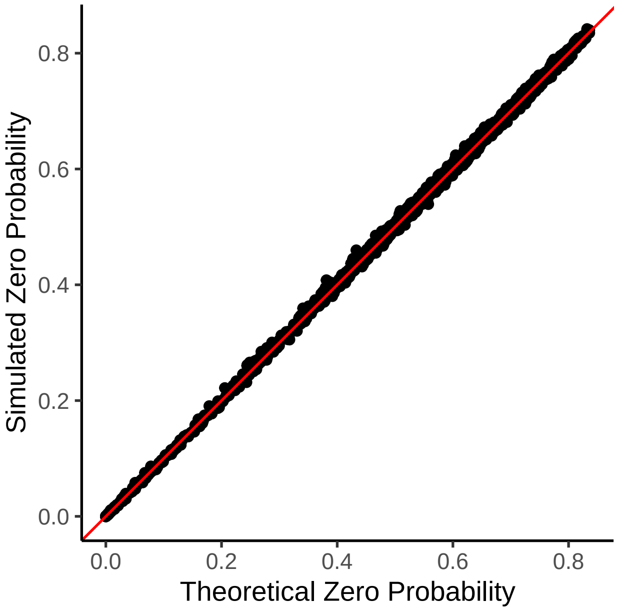


**Fig. S4**: Simulation results for zero read count probability vs. theoretical zero probability. Using the full generative process described for the molecular products and library sequencing, we observe that the probability that an observation is zero over multiple Monte Carlo simulations is a direct function of the initial rate parameter.

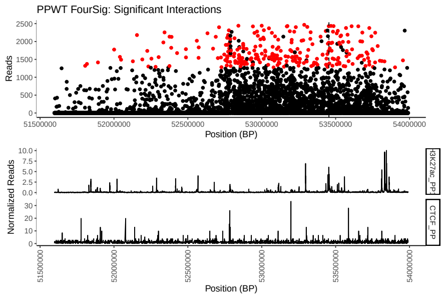


(a) ONECUT1 (FourSig)

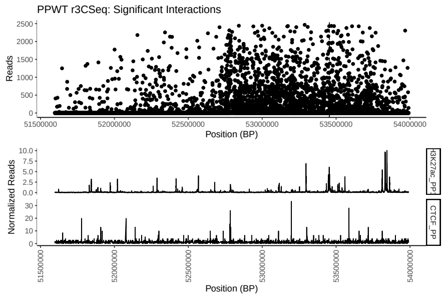


(b) ONECUT1 (r3Cseq)

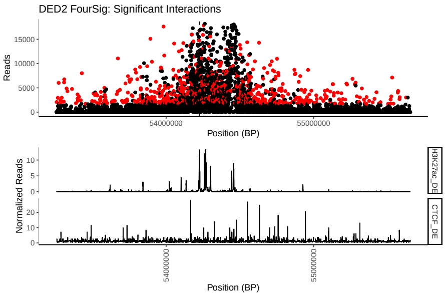


(c) SOX17 (FourSig)

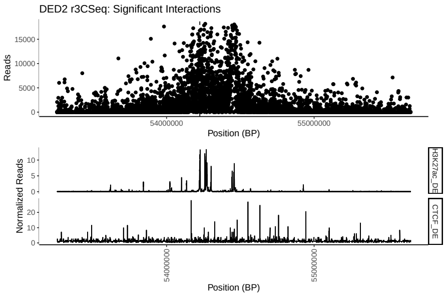


(d) SOX17 (r3Cseq)

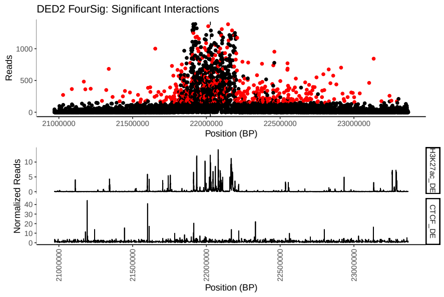


(e) GATA6 (FourSig)

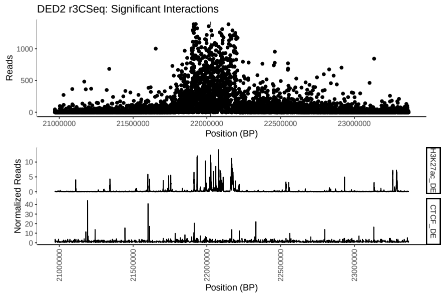


(f) GATA6 (r3Cseq)

**Fig. S5**: Significant interactions for *ONECUT1*, *SOX17*, and *GATA6* loci from FourSig and r3Cseq. The left column displays the pipeline results utilizing the FourSig methodology, while the right column shows the corresponding outputs generated via r3Cseq. All results are reported at an FDR of 0.1. Total counts across all replicates are shown. Bottom tracks show H3K27ac and CTCF respectively for the relevant conditions.

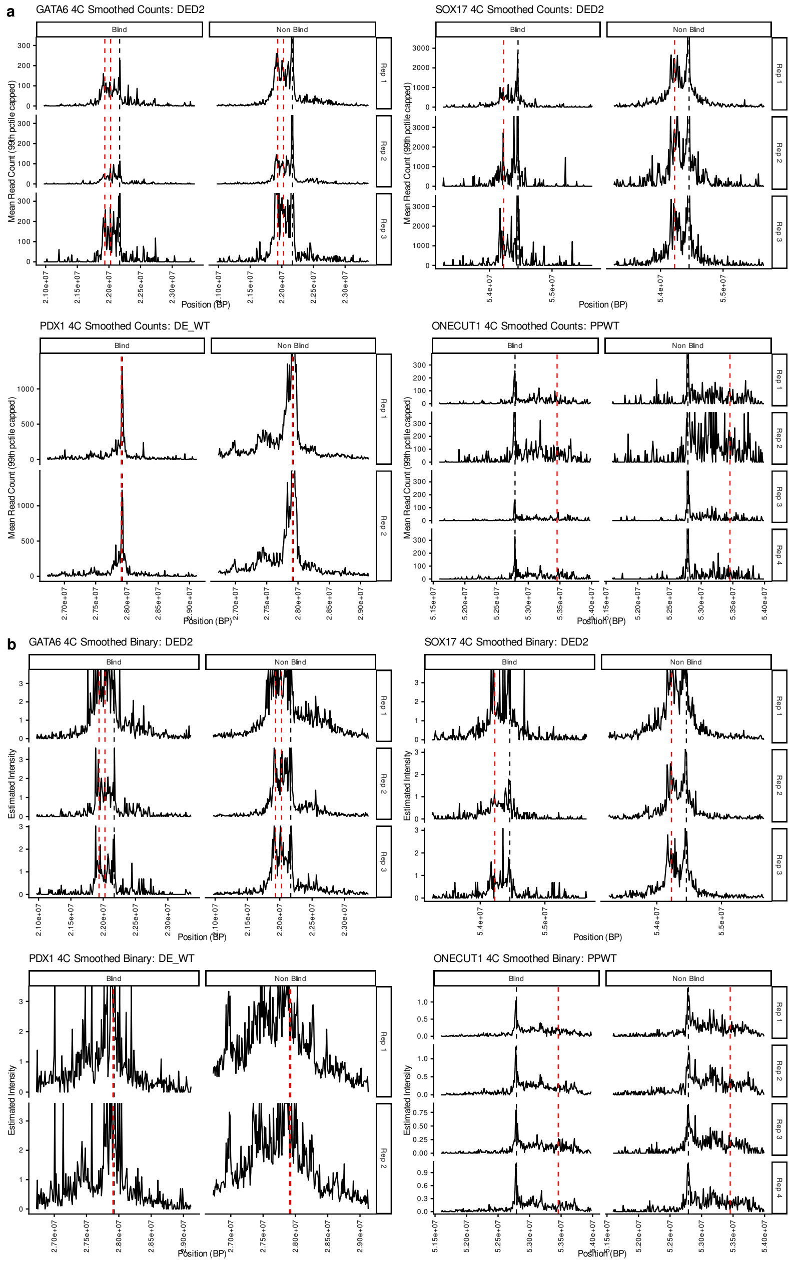


**Fig. S6**: For definitive endoderm samples, smoothed counts (a) and capture rates (b) were computed for *GATA6*,*SOX17*,*PDX1*,*ONECUT1* where the artifacts owing to PCR are removed by binarization.

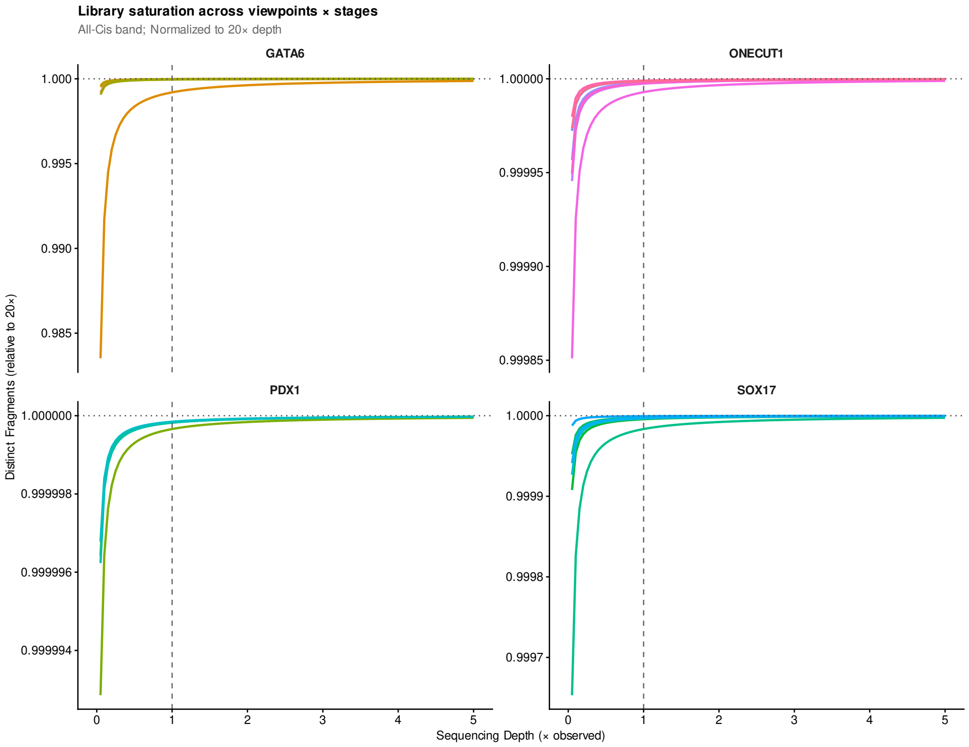


**Fig. S7**: For 4C-seq samples at *GATA6*,*SOX17*,*PDX1*,*ONECUT1* library complexity curves were obtained from PreseqR and scaled to the expected number of unique molecules that would have been observable at 20x the sequencing depth.

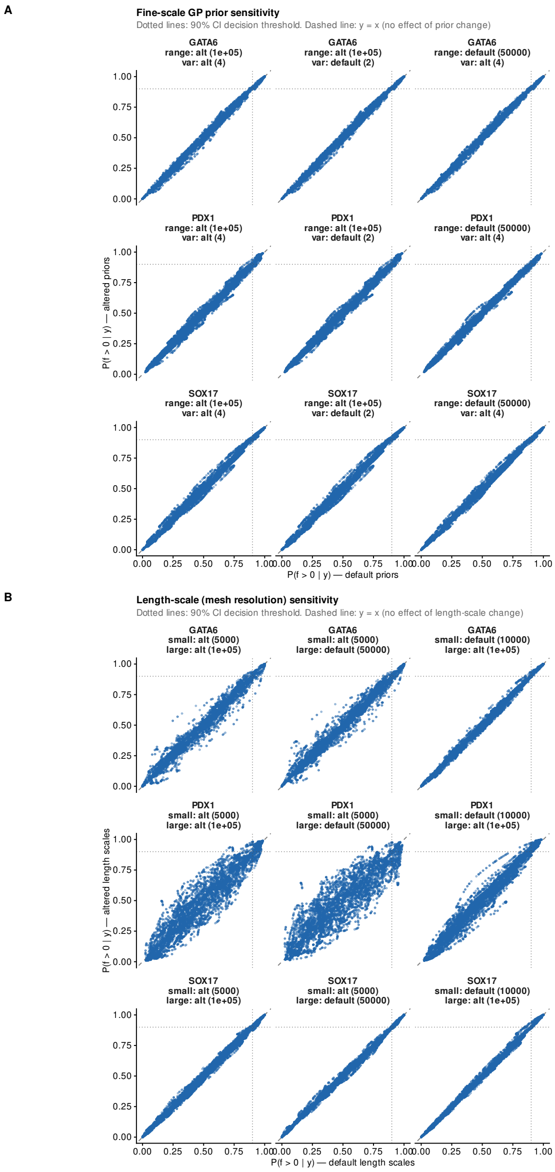


**Fig. S8**: For definitive endoderm samples, hyperparameters related to the Matern Gaussian process were varied. a) Across different viewpoints, the prior settings were varied from their default, and then the posterior probability of being a peak was compared to the default settings. b) Across different mesh grid spacings for both the fine and coarse scale terms, the posterior probabilities were compared to the default settings.

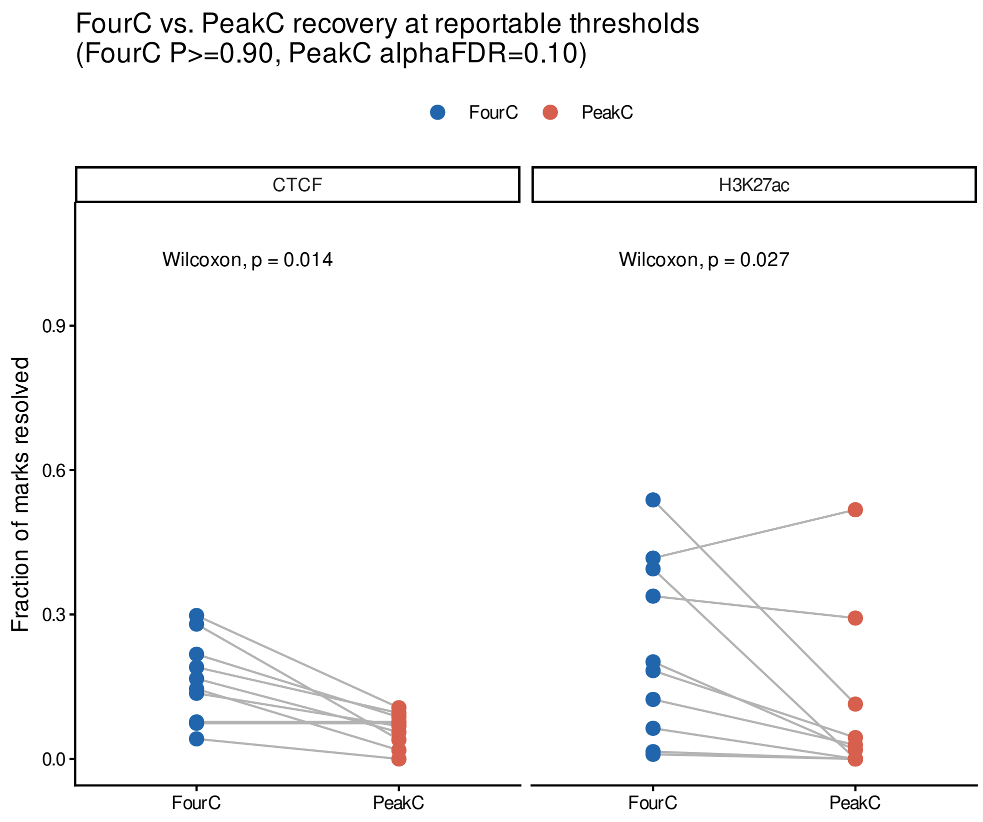


**Fig. S9**: Across different viewpoints, the recall of called CTCF or H3K27ac peaks in the vicinity of the viewpoints (±1.2 Mb) was assessed between both FourC and peakC, where in both cases, the differences are significant.

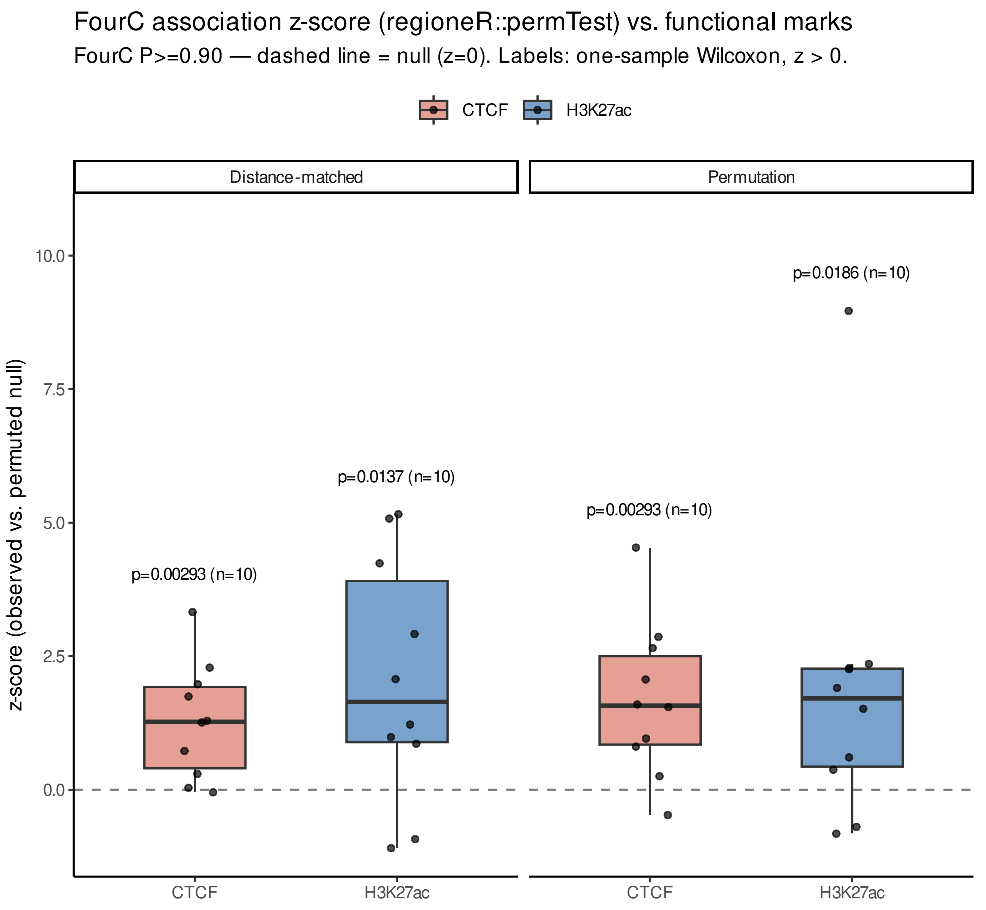


**Fig. S10**: Across different viewpoints, distance-matched blocked permutation controls were constructed to determine if FourC tends to enrich for the functional regions more often than expected by chance

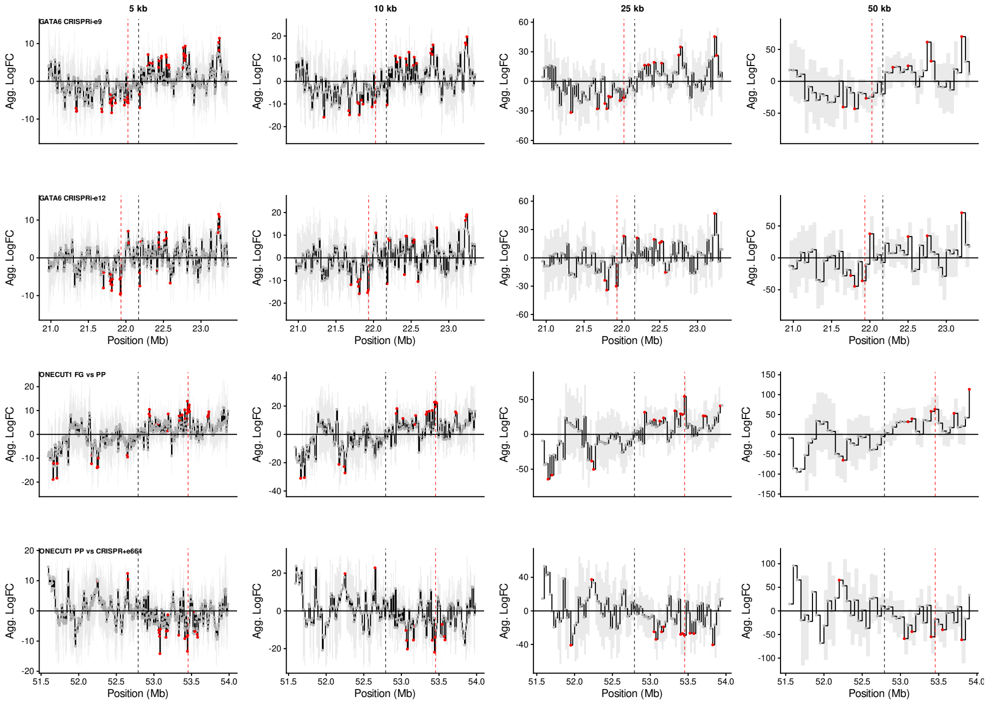


**Fig. S11**: Across the perturbation experiment cases, fragment-level FourC posteriors were binned at different sizes and then inference was performed on the aggregated posteriors.

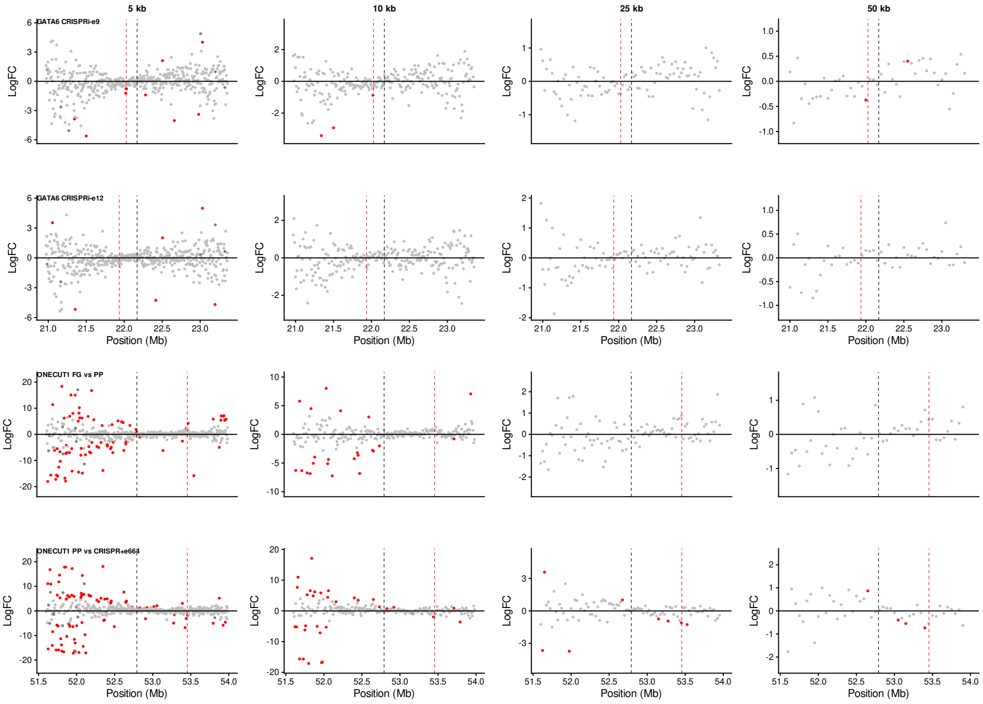


**Fig. S12**: Across the perturbation experiment cases, fragment-level data was first binned and then DESeq2 was applied to perform inferences on the fold changes between conditions.

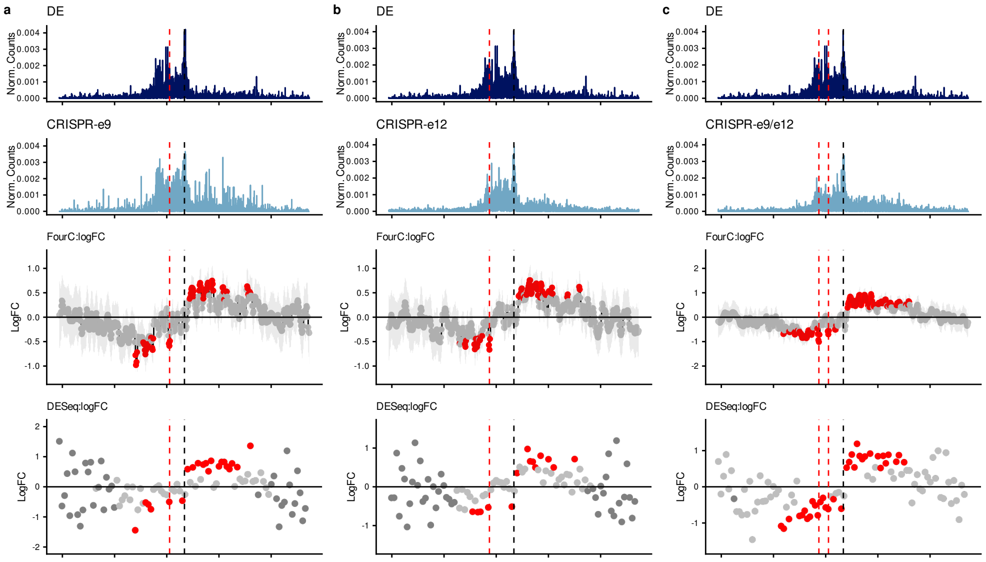


**Fig. S13**: CRISPR experiments at GATA6 produce strong changes in chromatin contact patterns, and lead to asymmetric changes whereby the TAD containing the gene tends to decrease in interaction frequency, while the adjacent region increases in interaction frequency. From top to bottom: raw data for the conditions being compared, FourC log fold change estimates, DESeq2 fold change estimates. Red dots indicate significant differences respectively if 90% central credible interval does not contain 0 or the FDR $\leq 0.1$

.

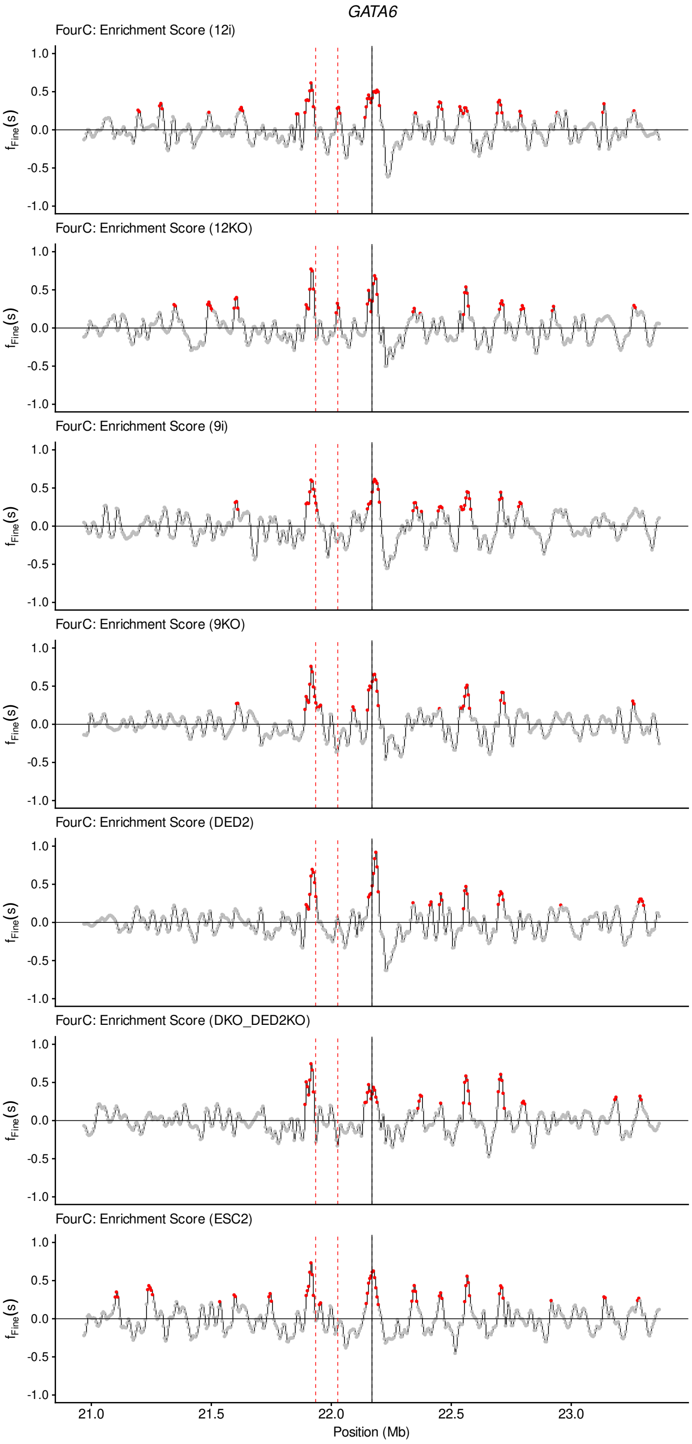


**Fig. S14**: Interactions called across *GATA6*

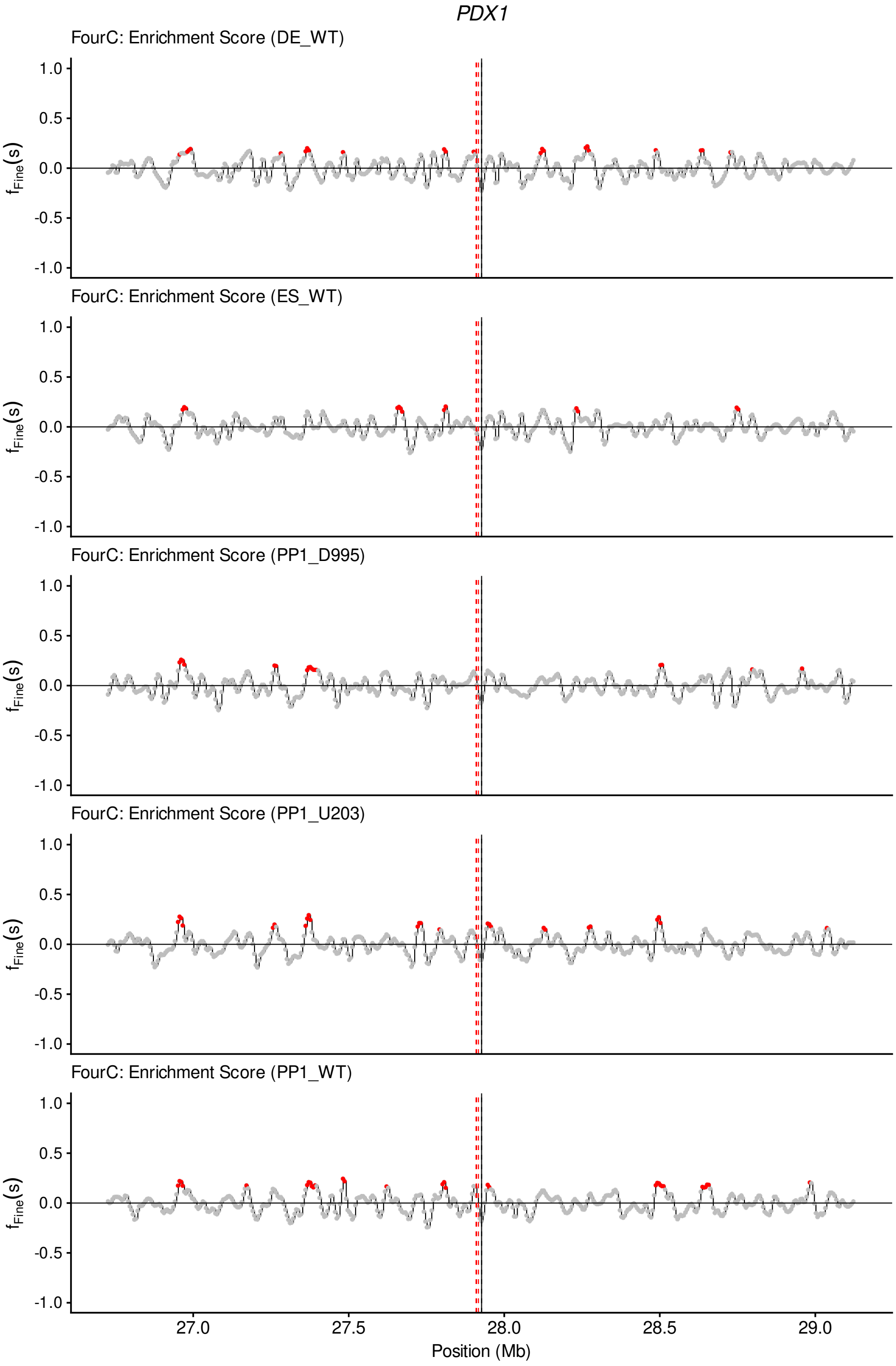


**Fig. S15**: Interactions called across *PDX1*

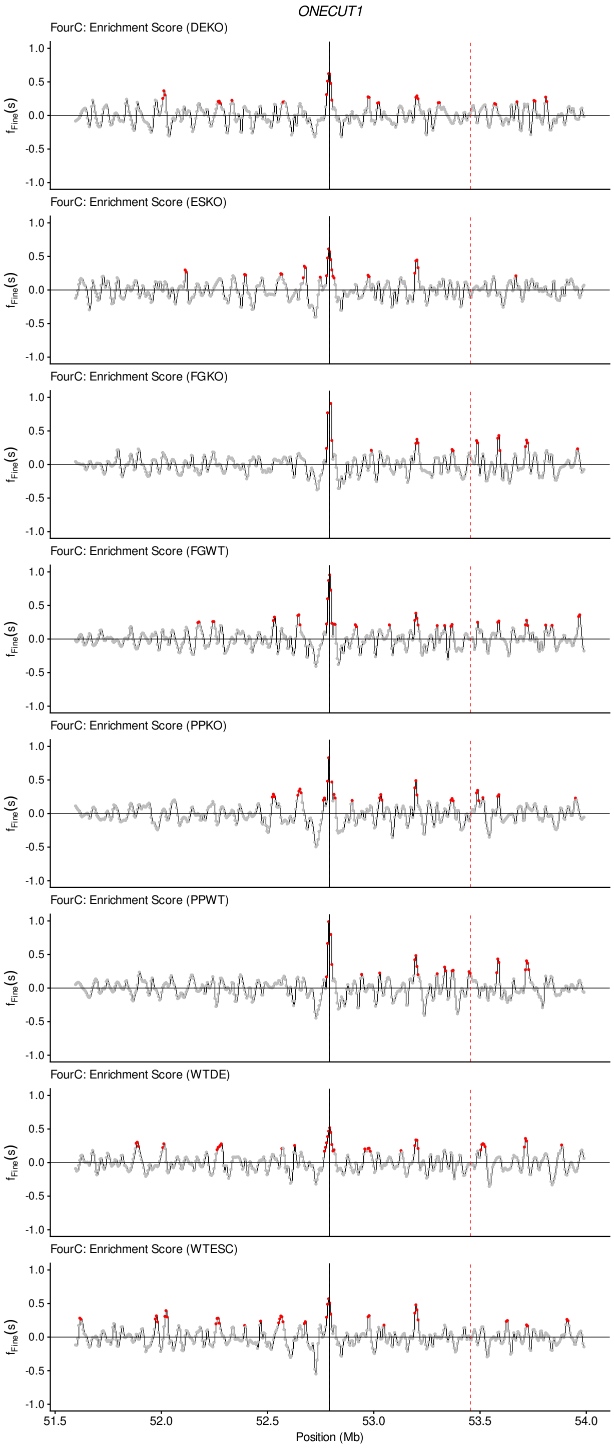


**Fig. S16**: Interactions called across *ONECUT1*

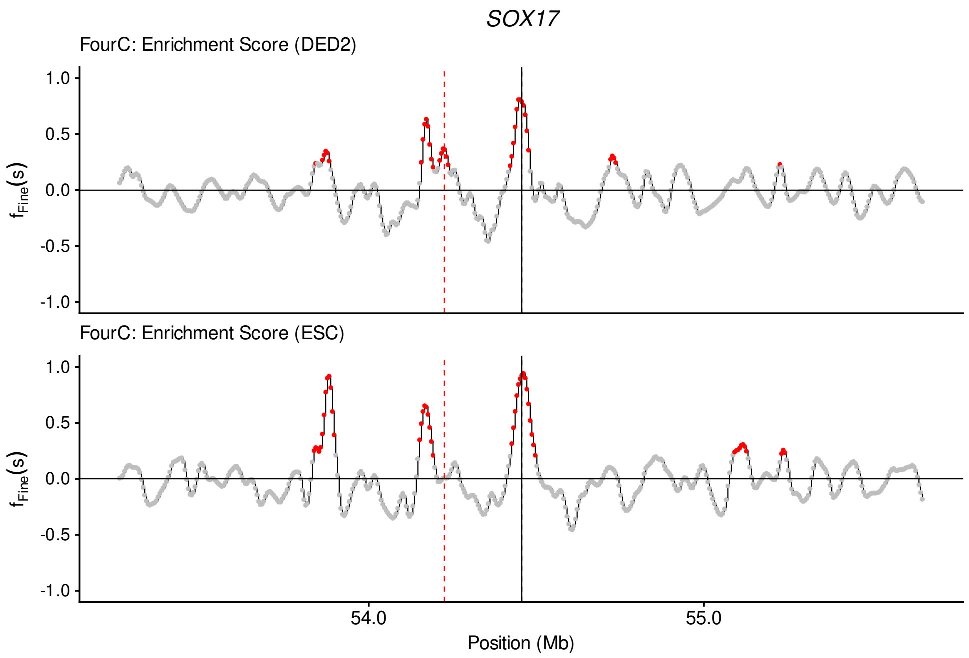


**Fig. S17**: Interactions called across *SOX17*

.
